# ISG15-dependent Activation of the RNA Sensor MDA5 and its Antagonism by the SARS-CoV-2 papain-like protease

**DOI:** 10.1101/2020.10.26.356048

**Authors:** GuanQun Liu, Jung-Hyun Lee, Zachary M. Parker, Dhiraj Acharya, Jessica J. Chiang, Michiel van Gent, William Riedl, Meredith E. Davis-Gardner, Effi Wies, Cindy Chiang, Michaela U. Gack

## Abstract

Activation of the RIG-I-like receptors, RIG-I and MDA5, establishes an antiviral state by upregulating interferon (IFN)-stimulated genes (ISGs). Among these is ISG15 whose mechanistic roles in innate immunity still remain enigmatic. Here we report that ISGylation is essential for antiviral IFN responses mediated by the viral RNA sensor MDA5. ISG15 conjugation to the caspase activation and recruitment domains of MDA5 promotes the formation of higher-order assemblies of MDA5 and thereby triggers activation of innate immunity against a range of viruses including coronaviruses, flaviviruses and picornaviruses. The ISG15-dependent activation of MDA5 is antagonized through direct de-ISGylation mediated by the papain-like protease (PLpro) of SARS-CoV-2, a recently emerged coronavirus that causes the COVID-19 pandemic. Our work demonstrates a crucial role for ISG15 in the MDA5-mediated antiviral response, and also identifies a novel immune evasion mechanism of SARS-CoV-2, which may be targeted for the development of new antivirals and vaccines to combat COVID-19.

## INTRODUCTION

Viral perturbation of host immune homeostasis is monitored by the innate immune system, which relies on receptors that sense pathogen-or danger-associated molecular patterns^1, 2, 3^. The RIG-I-like receptors (RLRs), RIG-I and MDA5 are pivotal for virus detection by surveying the cytoplasm for viral or host-derived immunostimulatory RNAs that harbor dsRNA structures and, in the case of RIG-I agonists, also a 5'-di-or tri-phosphate moiety^4^. Binding of RNA to the C-terminal domain (CTD) and helicase of RIG-I and MDA5 leads to their transition from an inactive state to a signaling-primed conformation that allows for the recruitment of several enzymes^5^. These enzymes modify RLRs at multiple domains and sites, and posttranslational modifications (PTMs) are particularly well studied for the N-terminal caspase activation and recruitment domains (CARDs), the signaling modules. PP1α/γ dephosphorylate specific CARD residues in RIG-I and MDA5^6^, which triggers further activation steps. In the case of RIG-I, dephosphorylation promotes K63-linked polyubiquitination of the CARDs by TRIM25 and other E3 ligases^7, 8^, which nucleates and stabilizes the oligomeric form of RIG-I, thereby enabling MAVS binding at mitochondria. Compared to those of RIG-I, the individual steps of MDA5 activation and critical PTMs involved are less well understood.

RLR activation induces the production of type I and III interferons (IFNs) which, in turn, propagate antiviral signaling by upregulating IFN-stimulated genes (ISGs)^9, 10^. Among the profoundly upregulated ISGs is ISG15, a ubiquitin-like protein. Similar to ubiquitin, ISG15 can be covalently conjugated to lysine (K) residues of target proteins, a PTM process termed ISGylation^11^. ISGylation is catalyzed by a chain of enzymatic reactions analogous to ubiquitination, involving an E1 activating enzyme (Ube1L), an E2 conjugating enzyme (UbcH8), and a handful of E3 ligases (for example, HERC5). Inversely, de-ISGylation is mediated by the cellular isopeptidase USP18^11^, and certain viruses also encode proteases that harbor de-ISGylase activities^12^. Besides covalent conjugation, ISG15 – like ubiquitin – can noncovalently bind to substrate proteins. Whereas ISG15 conjugation has been widely recognized to act antivirally^13^, unconjugated ISG15 serves a proviral role by promoting USP18-mediated suppression of type I IFN receptor (IFNAR) signaling^14, 15, 16^; this latter function of ISG15 is responsible for over-amplified ISG induction and fortified viral resistance in humans with inherited ISG15 deficiency. In contrast to ISG15’s role in dampening IFNAR signaling, the precise mechanism(s) of how ISGylation enhances immune responses to a wide range of viral pathogens are less well understood. Along these lines, although a broad repertoire of viral and cellular proteins has been shown to be targeted for ISGylation^13^ (of note, this usually represents co-translational modification of the nascent protein pool^17^), mechanisms of host protein ISGylation that could explain the broad antiviral restriction activity of ISG15 are currently unknown.

The causative agent of the ongoing COVID-19 pandemic, severe acute respiratory syndrome coronavirus 2 (SCoV2), belongs to the *Coronaviridae* family that contains several other human pathogens. Coronaviruses have an exceptional capability to suppress IFN-mediated antiviral responses, and low production of type I IFNs in SCoV-2-infected patients correlated with more severe disease outcome^18^. Among the coronaviral IFN antagonists is the papain-like protease (PLpro), which has deubiquitinating and de-ISGylating activities^19, 20^; however, the cellular substrates of the SCoV2 PLpro remain largely elusive.

Here we identify an essential role for ISGylation in MDA5 activation. We further show that SCoV2 PLpro interacts with MDA5 and antagonizes ISG15-dependent MDA5 activation via its de-ISGylase activity, unveiling that SCoV2 has already evolved to escape immune surveillance by MDA5.

## RESULTS

### MDA5, but not RIG-I, signaling requires ISG15

To identify PTMs of the CARDs of MDA5 that may regulate MDA5 activation, we subjected affinity-purified MDA5-2CARD fused to glutathione-*S*-transferase (GST-MDA5-2CARD), or GST alone, to liquid chromatography coupled with tandem mass spectrometry (LC-MS/MS) and found that specifically GST-MDA5-2CARD co-purified with ISG15, which appeared as two bands that migrated more slowly (by ~15 and 30 kDa) than unmodified GST-MDA5-2CARD (**Extended Data Fig. 1a**). Immunoblot (IB) analysis confirmed that GST-MDA5-2CARD is modified by ISG15 **(Extended Data Fig. 1b**). Since the CARDs are the signaling module of MDA5, we next determined the functional relevance of ISG15 for MDA5-induced signaling. While ectopic expression of FLAG-MDA5 in wild-type (WT) mouse embryonic fibroblasts (MEFs) induced IFN-β mRNA and protein as well as *Ccl5* transcripts in a dose-dependent manner, FLAG-MDA5 expression in *Isg15*−/− MEFs led to ablated antiviral gene and protein expression (**Fig. 1a and Extended Data Fig. 1c**). Similarly, antiviral gene expression induced by FLAG-MDA5 was strongly diminished in *ISG15* KO HeLa (human) cells compared to WT control cells (**Fig. 1b and Extended Data Fig. 1d**), ruling out a species-specific effect. In contrast to FLAG-MDA5, ectopically expressed FLAG-RIG-I induced comparable amounts of secreted IFN-β protein as well as *Ifnb1* and *Ccl5* transcripts in *Isg15*−/− and WT MEFs (**Fig. 1a and Extended Data Fig. 1c**). *IFNB1* transcripts and IFN-β protein production triggered by FLAG-RIG-I were slightly enhanced in *ISG15* KO HeLa cells as compared to WT control cells (**Fig. 1b and Extended Data Fig. 1d**), consistent with previous reports that ISGylation negatively impacts RIG-I signaling^21, 22^. These results suggest that ISG15 is required for MDA5, but not RIG-I, mediated signal transduction.

**Figure 1.**
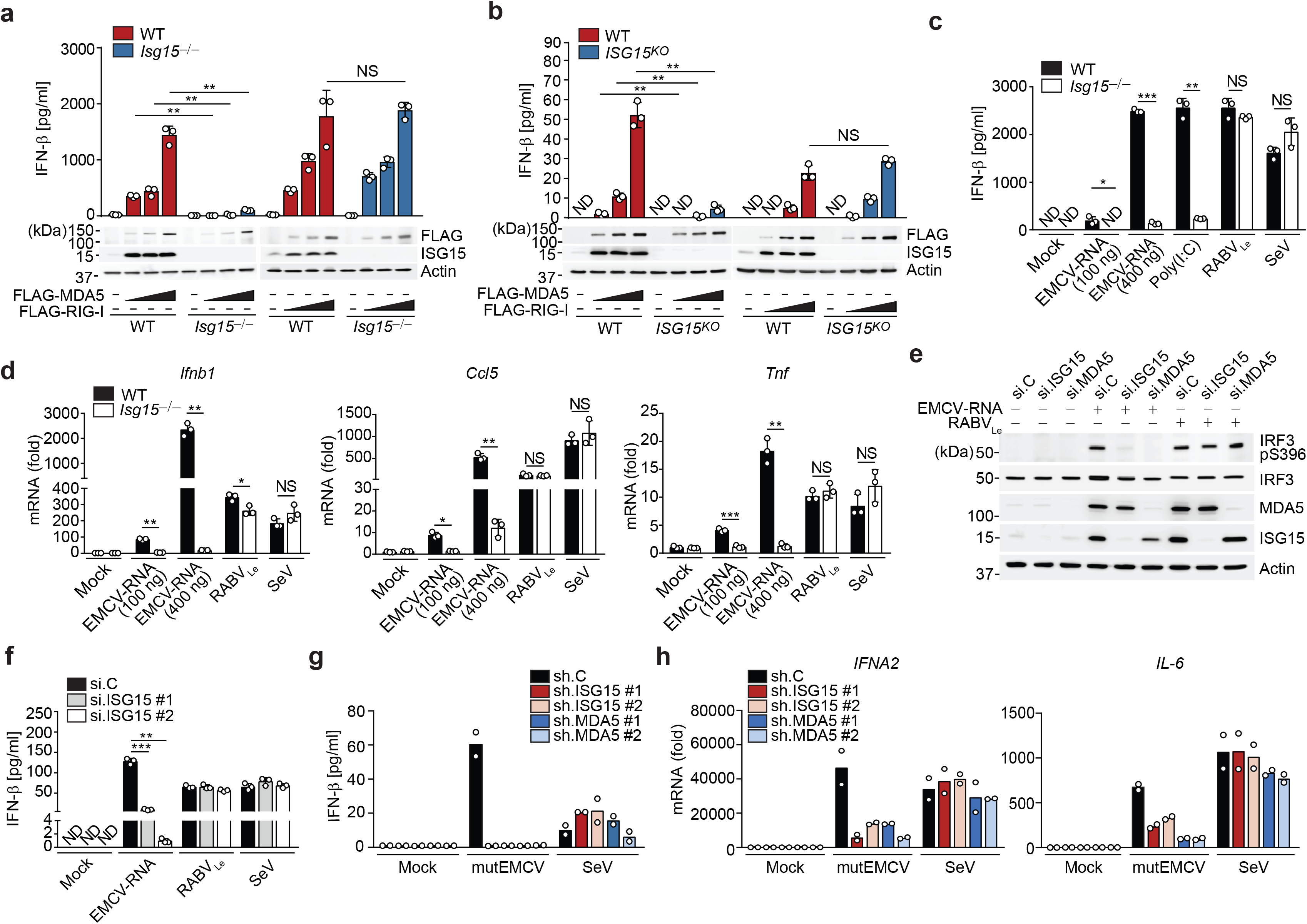
ISGylation is required for MDA5-, but not RIG-I signaling. **(a, b)** ELISA of IFN-β from supernatants of MEFs (WT or *Isg15*−/−) (a) and HeLa cells (WT or *ISG15* KO) (b) transiently transfected with increasing amounts of FLAG-tagged MDA5 or RIG-I for 40 h. Whole cell lysates (WCLs) were probed by immunoblotting (IB) with anti-ISG15, anti-FLAG, and anti-Actin (loading control). **(c)** ELISA of IFN-β from supernatants of WT or *Isg15*−/− MEFs that were mock-stimulated or transfected with EMCV-RNA (0.1 or 0.4 μg/mL), HMW-poly (I:C) (0.5 μg/mL), or RABVLe (1 pmol/mL), or infected with SeV (10 HAU/mL) for 24 h. **(d)** Quantitative RT-PCR (qRT-PCR) analysis of *IFNB1* and *CCL5* mRNA in WT and *Isg15*−/− MEFs stimulated as in (c). **(e)** IRF3 phosphorylation in the WCLs of NHLFs that were transfected with the indicated siRNAs for 30 h and then mock-stimulated or transfected with EMCV-RNA (0.4 μg/mL) or RABVLe (1 pmol/mL) for 6 h, assessed by IB with anti-pS396-IRF3 and anti-IRF3. **(f)** ELISA of IFN-β from supernatants of NHLFs that were transfected with the indicated siRNAs for 30 h and then mock-stimulated or transfected with EMCV-RNA (0.4 μg/mL) or RABVLe (1 pmol/mL), or infected with SeV (10 HAU/mL) for 16 h. **(g)** ELISA of IFN-β from the supernatants of PBMCs that were transduced for 40 h with the indicated shRNAs and then infected with mutEMCV (MOI 10) or SeV (200 HAU/mL) for 8 h. **(h)** qRT-PCR analysis of *IFNA2* and *IL-6* mRNA in PBMCs that were transduced and infected as in (g). Data are representative of at least two independent experiments (mean ± s.d. of *n* = 3 biological replicates in a, b, c, d, f and mean of *n* = 2 biological replicates in g and h). \**p* < 0.05, \*\**p* < 0.01, \*\*\**p* < 0.001 (unpaired Student’s *t*-test). ND, not detected; NS, not significant.

To substantiate a differential role of ISG15 in regulating MDA5 and RIG-I signaling, we tested the effect of *ISG15* gene deletion on the activation of endogenous MDA5 and RIG-I by their respective RNA ligands. IFN-β production as well as *IFNB1*, *CCL5*, and *TNF* gene expression induced by transfection of encephalomyocarditis virus (EMCV)-RNA or high-molecular-weight (HMW)-poly(I:C), both of which are predominantly sensed by MDA5, were profoundly attenuated in *Isg15*−/− MEF, *ISG15* KO HeLa, and *ISG15* KO HAP-1 (human) cells as compared to their respective control cells (**Fig. 1c,d and Extended Data Fig. 1e–g**). Importantly, the ablation of antiviral gene induction in response to EMCV-RNA or HMW-poly(I:C) in *ISG15* KO cells was not due to abrogated *MDA5* gene expression; on the contrary, mRNA expression of endogenous MDA5 was enhanced in *ISG15* KO cells as compared to WT cells (**Extended Data Fig. 1f,g)**. In contrast to stimulation with MDA5 agonists, stimulation of *Isg15*−/− MEFs and *ISG15* KO HeLa cells by transfection of rabies virus leader RNA (RABV_Le_) or by infection with Sendai virus (SeV, strain Cantell), both of which are specific RIG-I stimuli, led to IFN-β production and antiviral gene expression comparable to WT control cells (**Fig. 1c,d and Extended Data Fig. 1e**). To rule out potential clonal effects that could be associated with *ISG15* gene-deleted cells, we performed transient *ISG15* gene silencing in primary normal human lung fibroblasts (NHLFs) followed by stimulation of endogenous MDA5 and RIG-I with EMCV-RNA or RABV_Le_, respectively. siRNA-mediated silencing of *ISG15*, similarly to depletion of *MDA5,* led to a near-complete loss of phosphorylation of IFN-regulatory factor 3 (IRF3) – a hallmark of RLR signal activation – upon stimulation with EMCV-RNA, but not RABV_Le_ (**Fig. 1e**). In accord, knockdown of endogenous *ISG15* greatly diminished IFN-β production as well as *IFNB1* and *CCL5* gene expression in primary NHLFs transfected with EMCV-RNA, but not in cells transfected with RABVLe or infected with SeV (**Fig. 1f and Extended Data Fig. 1h**).

We next asked whether ISG15 is required for MDA5-mediated signaling also in immune cells. shRNA-mediated silencing of endogenous *ISG15* or *MDA5* in primary human peripheral blood mononuclear cells (PBMCs) substantially reduced IFN-β production and *IFNA2* and *IL-6* transcripts following infection with a recombinant mutant EMCV (mutEMCV) known to be deficient in MDA5 antagonism^23, 24^, as compared to infected PBMCs that were transduced with non-targeting control shRNA (**Fig. 1g,h and Extended Data Fig. 1i**). By contrast, *ISG15* or *MDA5* depletion did not affect the cytokine responses in PBMCs upon SeV infection. (**Fig. 1g,h and Extended Data Fig. 1i**). Collectively, these results show that ISG15 is essential for MDA5, but not RIG-I, mediated innate immune signaling.

### The MDA5 CARDs are ISGylated at K23 and K43

To corroborate our MS analysis that identified ISG15 modification of the MDA5-2CARD, we first tested whether also endogenous MDA5 is modified by ISG15. Anti-ISG15 immunoblot (IB) of immunoprecipitated endogenous MDA5 from primary NHLFs that were transfected with HMW-poly(I:C) or infected with the flaviviruses dengue (DENV) and Zika viruses (ZIKV) that are known to be sensed by MDA5 (together with RIG-I)^5^, showed robust ISGylation of MDA5 (**Fig. 2a**). Notably, endogenous MDA5 was also ISGylated in uninfected cells, although at low levels (**Extended Data Fig. 2a**), which is consistent with a previous report that showed that many host proteins are ISGylated at low levels also in normal (uninfected) conditions^17^. Moreover, in NHLFs that were treated with an anti-IFNAR2 antibody to block IFNAR-signaling-mediated ISG upregulation (*e.g*. *IFIT1* and *RSAD2*), silencing of *ISG15* or *MDA5* led to a comparable reduction of *IFNB1* gene expression in response to mutEMCV infection (**Extended Data Fig. 2b**). These results indicate that ISG15-dependent MDA5 signaling occurs even in the absence of IFNAR signaling, suggesting that basal ISGylation is sufficient for MDA5 activation.

**Figure 2.**
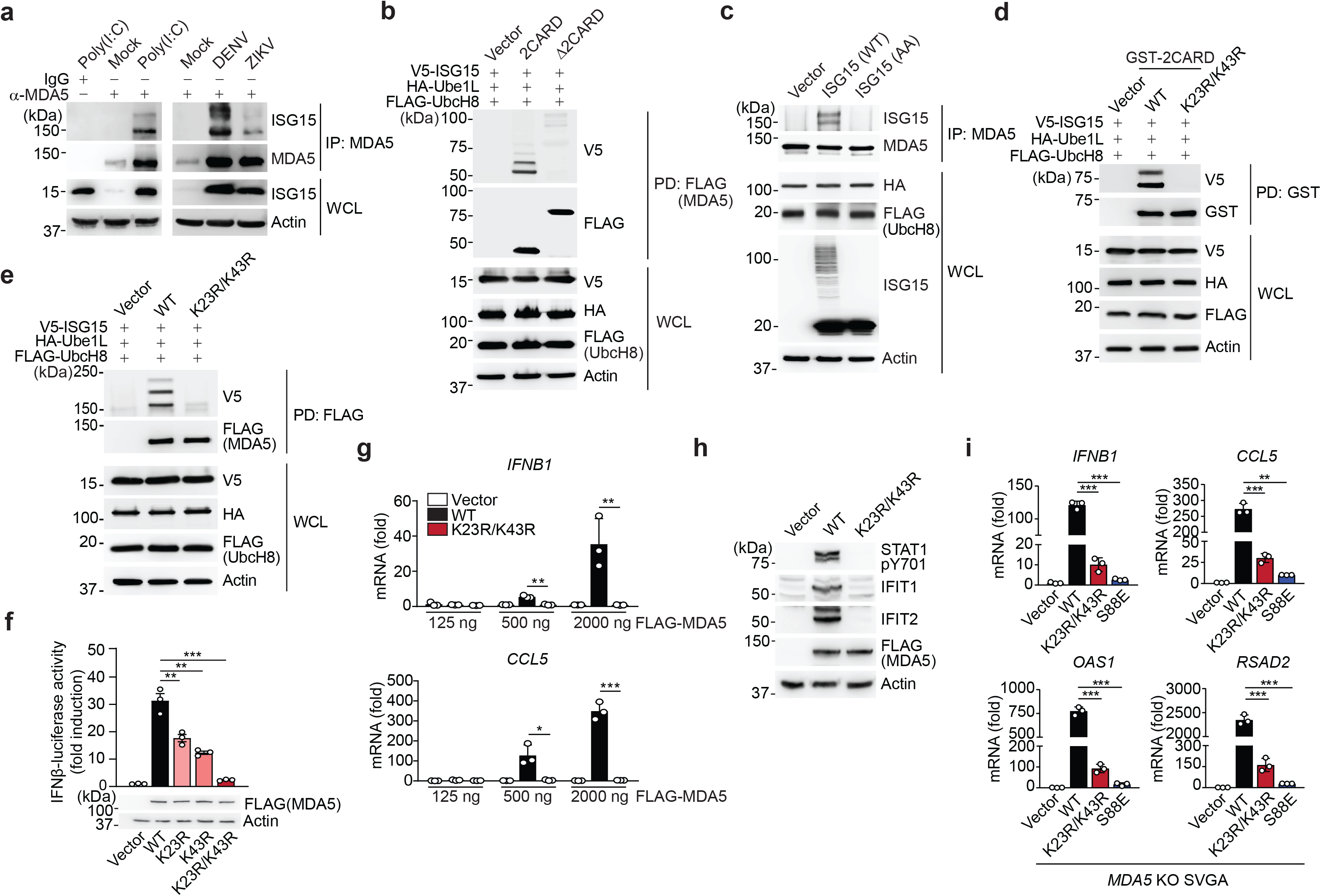
MDA5 activation requires ISGylation at K23 and K43. **(a)** Endogenous MDA5 ISGylation in NHLFs that were mock-treated, transfected with HMW-poly (I:C) (0.1 μg/mL) for 40 h (left), or infected with DENV or ZIKV (MOI 1 for each) for 48 h (right), determined by immunoprecipitation (IP) with anti-MDA5 (or an IgG isotype control) followed by IB with anti-ISG15 and anti-MDA5. WCLs were probed by IB with anti-ISG15 and anti-Actin (loading control). **(b)** ISGylation of FLAG-tagged MDA5-2CARD and MDA5ΔCARD in transiently transfected HEK293T cells that also expressed V5-ISG15, HA-Ube1L, and FLAG-UbcH8, assessed by FLAG pulldown (PD) and IB with anti-V5 and anti-FLAG forty hours after transfection. WCLs were probed by IB with anti-HA, anti-FLAG, anti-V5, and anti-Actin. **(c)** Endogenous MDA5 ISGylation in *ISG15* KO HeLa cells stably reconstituted with vector, WT ISG15 or ISG15-AA and co-transfected with HA-Ube1L and FLAG-UbcH8 after IFN-β treatment (1,000 U/mL) for 24 h, determined by IP with anti-MDA5 and IB with anti-ISG15 and anti-MDA5. **(d)** ISGylation of GST-MDA5-2CARD WT and K23R/K43R in HEK293T cells that were co-transfected with V5-ISG15, HA-Ube1L, and FLAG-UbcH8 for 24 h, determined by GST-PD and IB with anti-V5 and anti-GST. **(e)** ISGylation of FLAG-tagged MDA5 WT and K23R/K43R in HEK293T cells that were co-transfected with V5-ISG15, HA-Ube1L, and FLAG-UbcH8, determined by FLAG-PD and IB with anti-V5 and anti-FLAG. **(f)** IFN-β-luciferase reporter activity in HEK293T cells that were transfected for 40 h with vector, or FLAG-tagged MDA5 WT or mutants. Luciferase activity is presented as fold induction relative to the values for vector-transfected cells, set to 1. WCLs were probed by IB with anti-FLAG and anti-Actin. **(g)** qRT-PCR analysis of *IFNB1* and *CCL5* mRNA in HEK293T cells that were transiently transfected with either vector, or increasing amounts of FLAG-tagged MDA5 WT or K23R/K43R. **(h)** STAT1 phosphorylation and ISG (IFIT1 and 2) protein abundance in the WCLs of HEK293T cells that were transiently transfected with vector or FLAG-tagged MDA5 WT or K23R/K43R, determined by IB with anti-pY701-STAT1, anti-STAT1, anti-IFIT1, anti-IFIT2, anti-FLAG (expression control) and anti-Actin (loading control). **(i)** qRT-PCR analysis of *IFNB1*, *CCL5*, *OAS1*, and *RSAD2* mRNA in *MDA5* KO SVGAs that were reconstituted with either empty vector or FLAG-tagged MDA5 WT, K23R/K43R or S88E. Data are representative of at least two independent experiments (mean ± s.d. of *n* = 3 biological replicates in f, g, and i). \**p* < 0.05, \*\**p* < 0.01, \*\*\**p* < 0.001 (unpaired Student’s *t*-test). NS, not significant.

Biochemical analysis confirmed that the MDA5-2CARD, but not MDA5 Δ2CARD (containing helicase and CTD), is the primary site of MDA5 ISG15 modification (**Fig. 2b**). Of note, immunoblotting showed two major bands of ISGylation for MDA5-2CARD (**Fig. 2b**), which is consistent with our MS analysis (**Extended Data Fig. 1a**). Reconstitution of *ISG15* KO HeLa cells with either WT ISG15, or an unconjugatable mutant of ISG15 in which the two C-terminal glycines needed for conjugation were replaced with alanine (ISG15 AA), demonstrated covalent ISG15 conjugation of MDA5 (**Fig. 2c**).

Mutation of individual K residues in GST-MDA5-2CARD to arginine (R) revealed that single-site mutation of K23 and K43 noticeably reduced ISGylation (**Extended Data Fig. 2c**), while combined mutation of these two residues (K23R/K43R) led to a near-complete loss of ISGylation (**Fig. 2d**). Introduction of the K23R/K43R mutations into full-length FLAG-MDA5 also markedly diminished ISGylation (**Fig. 2e**), and the FLAG-MDA5 K23R/K43R mutant persisted in a hypo-ISGylated state over a 72-h time course of EMCV-RNA stimulation (**Extended Data Fig. 2d**). Of note, the residual ISGylation seen in MDA5 K23R/K43R is likely due to additional, minor sites in the CARD and/or Δ2CARD. To strengthen the concept that K23 and K43 are primarily modified by ISGylation and not other PTMs, we assessed the effect of the K23R/K43R mutation on MDA5 SUMOylation and ubiquitination^5^. The K23R/K43R mutation, which leads to a near-complete loss of MDA5 ISGylation, had no effect on MDA5 CARD SUMOylation; GST-MDA5-2CARD WT and the K23R/K43R mutant showed comparable SUMOylation levels (**Extended Data Fig. 2e)**. Furthermore, whereas GST-RIG-I-2CARD was robustly ubiquitinated (which primarily represents covalent K63-linked ubiquitination^7^), neither GST-MDA5-2CARD WT nor the K23R/K43R mutant showed detectable levels of ubiquitination under the same conditions (**Extended Data Fig. 2f**), which is in agreement with previous findings^7^. Taken together, these results indicate that the MDA5 CARDs undergo ISGylation at two major sites, K23 and K43.

### CARD ISGylation is required for MDA5 activation

To determine the relevance of CARD ISGylation in MDA5-mediated signaling, we first compared the ability of MDA5-2CARD WT and of the mutants K23R, K43R and K23R/K43R to activate the IFN-β promoter by luciferase reporter assay. Consistent with their reduced ISGylation levels (**Fig. 2d and Extended Data Fig. 2c**), MDA5-2CARD K23R and K43R single-site mutants showed partially reduced IFN-β promoter activation as compared to WT MDA5-2CARD, while the MDA5-2CARD K23R/K43R double mutant had a profoundly reduced signaling activity (**Extended Data Fig. 2g**). The decrease in signaling ability of the MDA5-2CARD K23R/K43R mutant was almost as strong as that of the phosphomimetic mutants S88E and S88D, which are inactive due to constitutive CARD ‘phosphorylation’ and thus served as positive controls^6^. In contrast, an MDA5-2CARD mutant in which K68, which is the lysine residue that is most proximal to K43 and K23, was substituted with arginine (K68R), showed comparable ISG15 conjugation and signaling competency to the WT 2CARD (**Extended Data Fig. 2c,g**). Consistent with the data obtained from the IFN-β luciferase assay, the MDA5-2CARD K23R/K43R mutant, in contrast to WT MDA5-2CARD, also failed to induce the dimerization of endogenous IRF3 (**Extended Data Fig. 2h**). Full-length FLAG-MDA5 K23R, K43R, or K23R/K43R double mutant, also showed reduced and near-abolished IFN-β promoter activating abilities, respectively, as compared to WT FLAG-MDA5 (**Fig. 2f**), strengthening that K23 and K43 are the ISGylation sites that are critical for MDA5 activation. Of note, the FLAG-MDA5 K23/K43R mutant showed a profound signaling defect even when expressed at high amounts. In contrast, WT FLAG-MDA5 induced *IFNB1* and *CCL5* transcript expression in a dose-dependent manner (**Fig. 2g)**. Consistent with these data, phosphorylation of STAT1, which is a hallmark of IFNAR-signal activation, as well as protein expression of IFIT1 and IFIT2 (both are ISGs) were highly induced in cells expressing WT MDA5, but not in cells expressing the K23R/K43R mutant (**Fig. 2h**). To rule out the possibility of a confounding effect by endogenous MDA5 on signaling induced by our ectopically-expressed MDA5 mutants, we tested their signal-transducing activities in human astrocytes in which the *MDA5* gene expression was ablated using CRISPR-Cas9 technology (*MDA5* KO SVGAs) (**Fig. 2i and Extended Data Fig. 2i**). Complementation of *MDA5* KO SVGA cells with the K23R/K43R mutant led to greatly diminished *IFNB1*, *CCL5*, and ISG (*OAS1* and *RSAD2*) transcript induction compared to cells expressing WT MDA5. Control cells reconstituted with the signaling-defective MDA5 S88E mutant also showed strongly reduced antiviral gene induction (**Fig. 2i**). These results demonstrate that ISGylation at K23 and K43 in the CARDs is essential for MDA5-mediated antiviral cytokine responses.

### Dephosphorylation by PP1 regulates MDA5 ISGylation

Like RIG-I, MDA5 is phosphorylated within the CARDs in uninfected cells, which prevents auto-activation; in contrast, dephosphorylation of RIG-I (at S8 and T170) and MDA5 (at S88) by PP1α/γ is crucial for unleashing RLRs from their signaling-repressed states^6, 25, 26, 27^. In the case of RIG-I, dephosphorylation allows K63-linked ubiquitination of the CARDs, which then promotes RIG-I multimerization and antiviral signaling^5^. The details of how CARD dephosphorylation triggers MDA5 activation have remained elusive, and therefore we tested whether dephosphorylation regulates MDA5 ISGylation. We found that silencing of endogenous PP1α or PP1γ strongly diminished MDA5-2CARD ISGylation (**Extended Data Fig. 3a**). Furthermore, the phosphomimetic MDA5-2CARD mutants S88E and S88D had markedly reduced ISGylation, whereas the ‘phospho-null’ S88A mutant showed stronger ISGylation than WT MDA5-2CARD (**Extended data Fig. 3b**). Conversely, the ISGylation-null mutant of MDA5, K23R/K43R, had comparable S88 phosphorylation levels (**Extended Data Fig. 3c**). Together, these data suggested that MDA5 dephosphorylation at S88 precedes CARD ISGylation.

**Figure 3.**
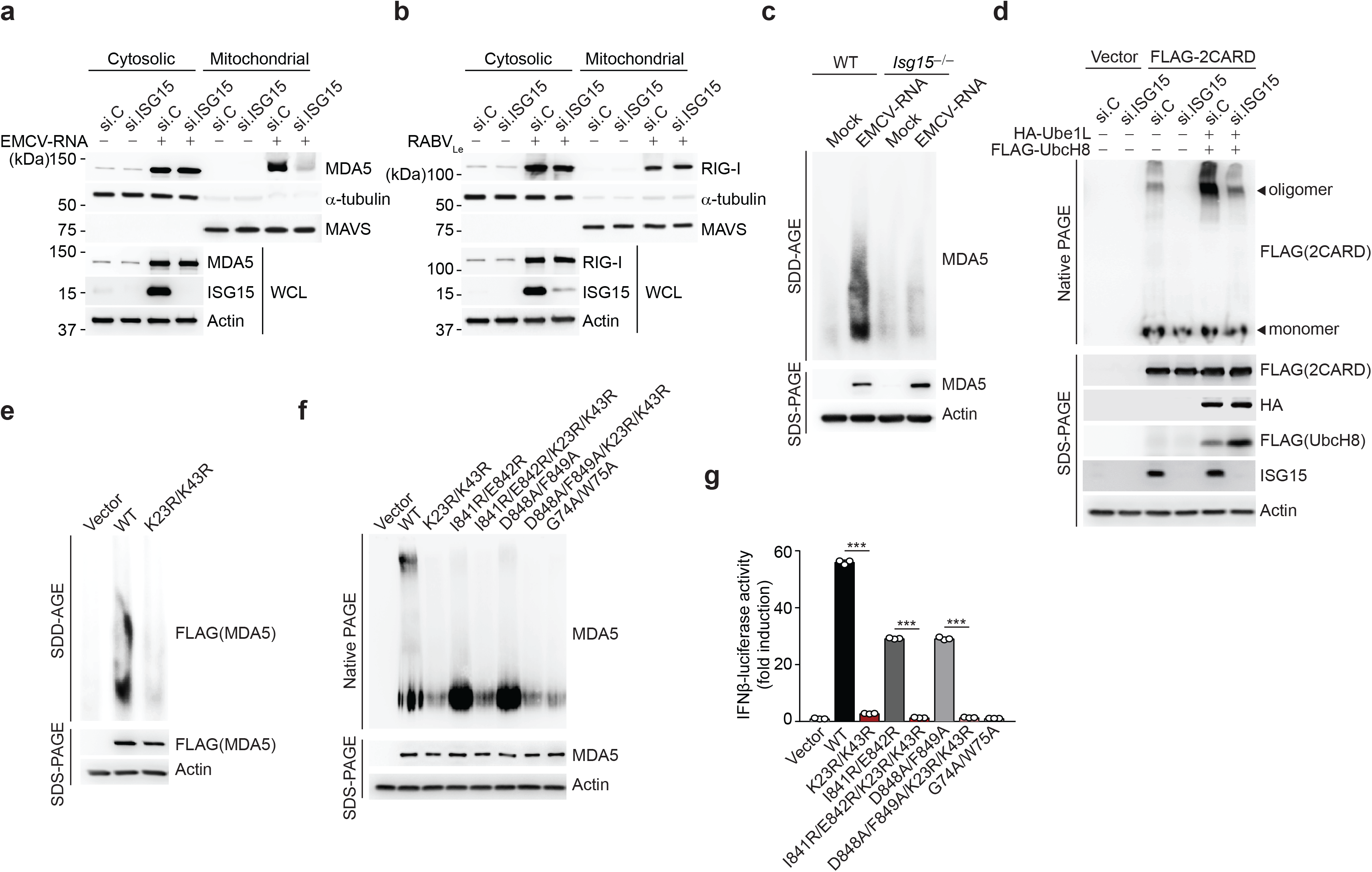
CARD ISGylation is essential for formation of higher-order MDA5 assemblies. (**a**,**b)** Cytosol-mitochondria fractionation of WCLs from NHLFs that were transfected for 30 h with non-targeting control siRNA (si.C) or ISG15-specific siRNA (si.ISG15) and then mock-treated or transfected with EMCV-RNA (0.4 μg/mL) (a) or RABVLe (1 pmol/mL) (b) for 16 h. IB was performed with anti-MDA5 (a), anti-RIG-I (b), anti-ISG15 and anti-Actin (a, b). α-Tubulin and MAVS served as purity markers for the cytosolic and mitochondrial fraction, respectively (a, b). **(c)** Endogenous MDA5 oligomerization in WT and *Isg15*−/− MEFs that were transfected with EMCV-RNA (0.5 μg/mL) for 16 h, assessed by SDD-AGE and IB with anti-MDA5. WCLs were further analyzed by SDS-PAGE and probed by IB with anti-MDA5 and anti-Actin. **(d)** Oligomerization of FLAG-MDA5-2CARD in HEK293T cells that were transfected with the indicated siRNAs together with or without HA-Ube1L and FLAG-UbcH8 for 48 h, determined by native PAGE and IB with anti-FLAG. WCLs were further analyzed by SDS-PAGE and probed by IB with anti-FLAG, anti-HA, anti-ISG15, and anti-Actin. **(e)** Oligomerization of FLAG-MDA5 WT and K23R/K43R in transiently transfected *MDA5* KO HEK293 cells, assessed by SDD-AGE and IB with anti-FLAG. WCLs were further analyzed by SDS-PAGE and IB with anti-FLAG and anti-Actin. **(f)** Oligomerization of FLAG-tagged MDA5 WT and mutants in transiently transfected *MDA5* KO HEK293 cells, assessed by native PAGE and IB with anti-MDA5. WCLs were further analyzed by SDS-PAGE and probed by IB with anti-MDA5 and anti-Actin. **(g)** IFN-β-luciferase reporter activity in *MDA5* KO HEK293 cells that were transfected for 24 h with either empty vector, or FLAG-tagged MDA5 WT or mutants. Luciferase activity is presented as fold induction relative to the values for vector-transfected cells, set to 1. Data are representative of at least two independent experiments (mean ± s.d. of *n* = 3 biological replicates in f). \*\*\**p* < 0.001 (unpaired Student’s *t*-test).

We next made use of the V protein of measles virus (MeV-V) of the *Paramyxoviridae* family, which is known to antagonize MDA5 S88 dephosphorylation through direct antagonism of PP1α/γ^28^. Ectopic expression of MeV-V enhanced the S88 phosphorylation (indicative of inhibition of S88 dephosphorylation) of GST-MDA5-2CARD or FLAG-MDA5 in a dose-dependent manner, as previously shown^28^. The enhancement of S88 phosphorylation by MeV-V correlated with a gradual decline in ISGylation (**Extended data Fig. 3d,e**). In contrast to WT MeV-V, a C-terminally truncated mutant of MeV-V (MeV-V Δtail) which has abolished PP1-binding and MDA5-dephosphorylation antagonism^28^, exhibited little effect on MDA5-2CARD ISGylation (**Extended data Fig. 3f**), strengthening that the inhibition of MDA5-2CARD ISGylation is primarily due to PP1 inhibition, and not other antagonistic effects, by the MeV-V protein. The V proteins from Nipah and Hendra viruses (NiV-V and HeV-V) also strongly enhanced MDA5 S88 phosphorylation (**Extended data Fig. 3g,h**), and correspondingly, dampened MDA5 ISGylation (**Extended data Fig. 3h**), suggesting that certain paramyxoviral V proteins inhibit MDA5 ISGylation through manipulation of S88 phosphorylation, although the precise mechanisms for individual V proteins remain to be determined. Taken together, these data suggest that the ISGylation of MDA5-2CARD is dependent on dephosphorylation at S88.

### ISGylation promotes higher-order MDA5 assemblies

The activation of RLRs is a multi-step process that includes RNA binding, RLR oligomerization, and their translocation from the cytosol to mitochondria and mitochondria-associated membranes for an interaction with MAVS^5^. To elucidate the mechanism by which ISGylation impacts MDA5 activity, we first examined whether ISGylation affects the ability of MDA5 to bind dsRNA. Endogenous MDA5 purified from WT or *Isg15*−/− MEFs interacted equally well with HMW-poly(I:C) *in vitro* (**Extended Data Fig. 4a**). Moreover, MDA5 WT and the K23R/K43R mutant showed comparable binding to HMW-poly(I:C), indicating that ISGylation does not affect the RNA-binding ability of MDA5 (**Extended Data Fig. 4b**). Next, we monitored the translocation of endogenous MDA5 from the cytosol to mitochondria in cells that were either depleted of *ISG15* using siRNA, or transfected with nontargeting control siRNA (si.C). EMCV-RNA-induced cytosol-to-mitochondria translocation of MDA5 was abolished in *ISG15*-silenced cells, whereas si.C-transfected cells showed efficient translocation (**Fig. 3a**). In contrast, the translocation of endogenous RIG-I from the cytosol to mitochondria induced by RABV_Le_ transfection was efficient in both *ISG15*-depleted and si.C-transfected cells (**Fig. 3b**). These data indicated that ISGylation influences MDA5 activation at the level of translocation, or a step upstream of it. Since the cytosol-to-mitochondria translocation of MDA5 has been shown to require an interaction with the chaperon protein 14-3-3η^29^, we assessed 14-3-3η-binding of WT MDA5 and its mutants. Pulldown assay showed that the ability of the MDA5 K23R/K43R mutant to bind 14-3-3η was similar to that of WT MDA5 or the K68R mutant (**Extended Data Fig. 4c**). However, whereas EMCV-RNA stimulation effectively induced the oligomerization of endogenous MDA5 in WT MEFs, the formation of MDA5 oligomers was ablated in MEFs that were deficient in *ISG15* (**Fig. 3c**). Consistent with these data, silencing of *ISG15* in human (293T) cells abolished the oligomerization of FLAG-MDA5-2CARD (**Fig. 3d**). Furthermore, co-expression of the ISGylation machinery components, Ube1L and UbcH8, strongly enhanced MDA5-2CARD oligomerization in si.C-transfected cells, but not in *ISG15*-depleted cells (**Fig. 3d**), indicating that ISGylation is required for MDA5 oligomer formation. In support of this concept, full-length FLAG-MDA5 K23R/K43R mutant showed near-abolished oligomerization, while WT MDA5 oligomerized efficiently (**Fig. 3e**). We also compared the effect of the K23R/K43R mutation with that of a panel of previously characterized oligomerization-disruptive mutations on the ability of MDA5 to oligomerize and signal downstream (**Fig. 3f,g**). These mutations localize either to the interface between MDA5 monomers (I841R/E842R and D848A/F849A) and impede RNA-binding-mediated MDA5 filamentation^30, 31^, or they localize to the CARDs (G74A/W75A) and disrupt 2CARD oligomerization^30^. In contrast to WT MDA5, the K23R/K43R mutant, similarly to the G74A/W75A mutant, showed deficient oligomerization and, consistent with this, abolished IFN-β promoter-activating ability (**Fig. 3f,g)**. Introduction of K23R/K43R into the I841R/E842R or D848A/F849A background, either of which by itself decreased MDA5 oligomerization and signaling, also abolished the formation of MDA5 oligomers and IFN-β promoter activation (**Fig. 3f,g**), suggesting a dominant role for CARD ISGylation in the formation of higher-order MDA5 assemblies. Since LGP2, the third member of the RLR family, has been shown to facilitate MDA5 nucleation on dsRNA and thereby MDA5 oligomerization^32, 33^, we tested the binding of LGP2 to MDA5 WT or K23R/K43R by Co-IP. MDA5 K23R/K43R mutant interacted with LGP2 as efficiently as WT MDA5 (**Extended Data Fig. 4d**), strengthening that MDA5 CARD ISGylation promotes MDA5 oligomerization independently of RNA-binding-mediated filamentation. Collectively, these results establish that ISGylation of the MDA5 CARDs potentiates MDA5 signaling by facilitating CARD oligomerization and formation of higher-order MDA5 assemblies.

**Figure 4.**
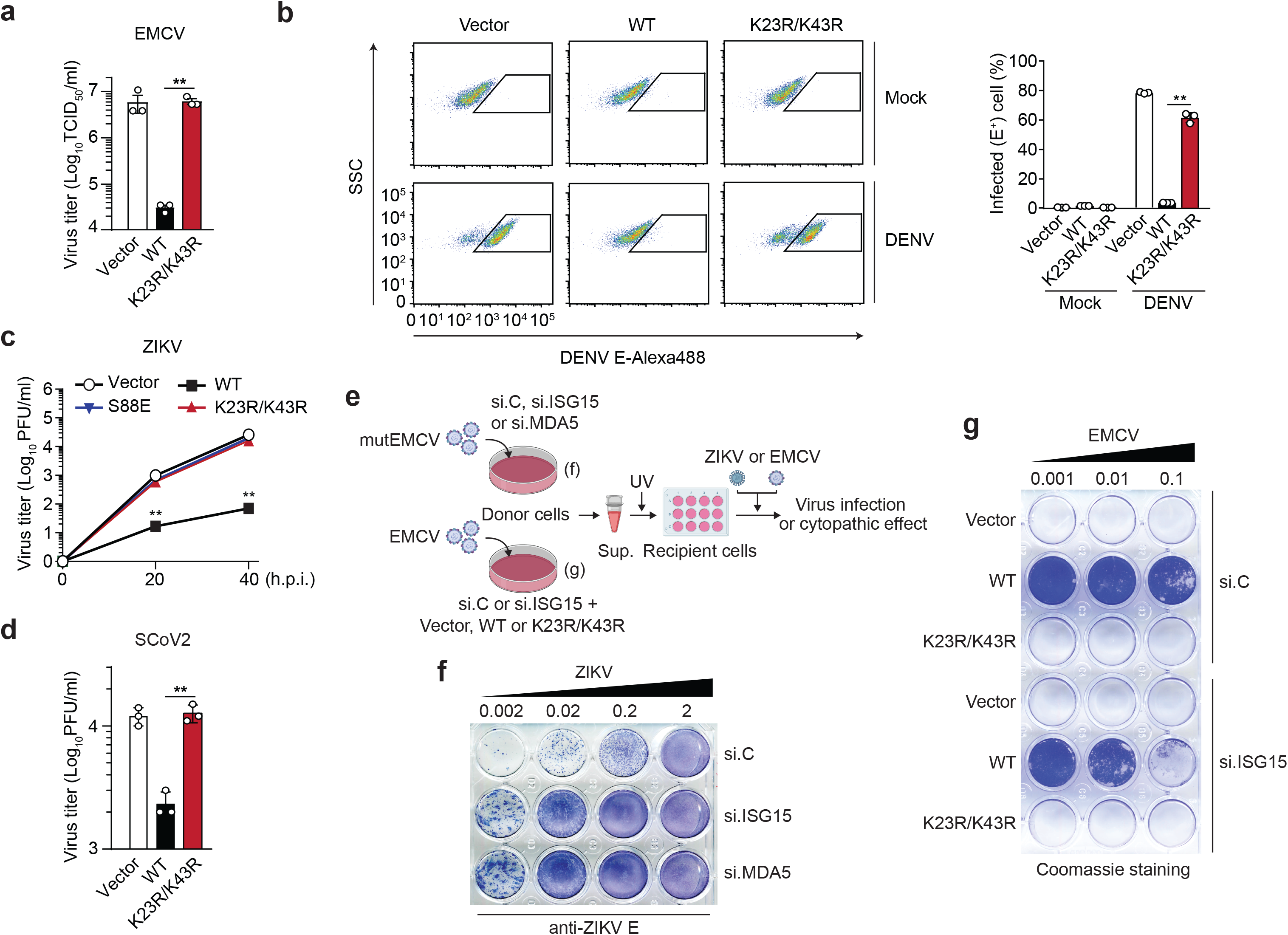
ISGylation is required for viral restriction by MDA5. **(a)** EMCV titers in the supernatant of HEK293T cells that were transiently transfected for 40 h with either empty vector, or FLAG-tagged MDA5 WT or K23/K43R and then infected with EMCV (MOI 0.001) for 24 h, determined by TCID50 assay. **(b)** Percentage of DENV-infected *MDA5* KO HEK293 cells that were transiently transfected for 24 h with either empty vector or FLAG-tagged MDA5 WT or K23R/K43R and then mock-treated or infected with DENV (MOI 5) for 48 h, assessed by FACS using an anti-flavivirus E (4G2) antibody. SSC, side scatter. **(c)** ZIKV titers in the supernatant of *MDA5* KO SVGAs that were transiently transfected for 30 h with either empty vector, or FLAG-tagged MDA5 WT, K23R/K43R, or S88E and then infected with ZIKV (MOI 0.1) for the indicated times, determined by plaque assay. **(d)** SCoV2 titers in the supernatant of HEK293T-hACE2 cells that were transiently transfected for 24 h with either empty vector, or FLAG-tagged MDA5 WT or K23/K43R and then infected with SCoV2 (MOI 0.5) for 24 h, determined by plaque assay. **(e)** Schematic of the experimental approach to decouple the role of ISG15 in MDA5-mediated IFN induction from its role in dampening IFNAR signaling. **(f)** NHLF ‘donor’ cells were transfected for 40 h with the indicated siRNAs and then infected with mutEMCV (MOI 0.1) for 16 h. Cell culture supernatants were UV-inactivated and transferred onto Vero ‘recipient’ cells for 24 h, followed by infection of cells with ZIKV (MOI 0.002 to 2) for 72 h. ZIKV-positive cells were determined by immunostaining with anti-flavivirus E (4G2) antibody and visualized using the KPL TrueBlue peroxidase substrate. **(g)** *RIG-I* KO HEK293 ‘donor’ cells were transfected for 24 h with si.C or si.ISG15 and subsequently transfected with either empty vector or FLAG-tagged MDA5 WT or K23R/K43R for 24 h, followed by EMCV infection (MOI 0.001) for 16 h. UV-inactivated culture supernatants were transferred onto Vero ‘recipient’ cells for 24 h, followed by infection with EMCV (MOI 0.001 to 0.1) for 40 h. EMCV-induced cytopathic effects were visualized by Coomassie blue staining. Data are representative of at least two independent experiments (mean ± s.d. of *n* = 3 biological replicates in a, b, c). \*\**p* < 0.01 (unpaired Student’s *t*-test).

### ISGylation-dependent MDA5 signaling restricts virus replication

We next assessed whether ISGylation of MDA5 is required for its ability to restrict virus replication. Ectopic expression of FLAG-MDA5 WT, but not of the K23R/K43R mutant, potently (by ~2-log) inhibited the replication of EMCV, which is sensed by MDA5 (**Fig. 4a**). Similarly, *MDA5* KO HEK293 cells reconstituted with WT MDA5, but not cells complemented with the K23R/K43R mutant, effectively restricted DENV replication (**Fig. 4b**). We also reconstituted *MDA5* KO astrocyte SVGAs, a physiologically relevant cell type for ZIKV infection, with either vector, or MDA5 WT or K23R/K43R and then assessed ZIKV replication over a 40-hour time course. ZIKV replication was attenuated by ~100-fold in cells reconstituted with WT MDA5 as compared to vector-transfected cells. In contrast, cells complemented with MDA5 K23R/K43R did not restrict ZIKV growth, similarly to the signaling-defective S88E mutant, which served as an additional control (**Fig. 4c**). Similarly, ectopic expression of WT MDA5 restricted the replication of SARS-CoV-2 (SCoV2), a recently emergent coronavirus that is responsible for the ongoing COVID-19 pandemic. In contrast, MDA5 K23R/K43R did not inhibit SCoV2 replication (**Fig. 4d**).

To further substantiate that MDA5-mediated virus restriction is dependent on MDA5 ISGylation, we determined the effect of *ISG15* silencing on the ability of FLAG-MDA5 WT or K23R/K43R to inhibit EMCV replication. While the K23R/K43R mutant failed to suppress EMCV replication regardless of *ISG15* silencing, WT MDA5 effectively restricted EMCV replication in si.C-transfected cells, and unexpectedly, also in *ISG15* knockdown cells (**Extended Data Fig. 5a**). In an exploration of the underlying mechanism of these unexpected results, we found that the EMCV-infected cells that expressed WT MDA5 had markedly enhanced levels of ISG protein expression (*i.e*. IFIT1, IFIT2, RSAD2, and ISG20) when *ISG15* was silenced as compared to infected cells transfected with the control siRNA (**Extended Data Fig. 5b**). Similarly, elevated ISG transcript and protein expression was observed in *ISG15*-deficient cells that were transfected with EMCV-RNA or infected with mutEMCV, despite the abrogation of IFN-β induction (**Extended Data Fig. 5c,d**). In contrast, silencing of endogenous MDA5 abrogated both IFN-β production and ISG protein expression, as expected (**Extended Data Fig. 5d**). We noticed that the protein abundance of USP18, a deubiquitinating enzyme that negatively regulates IFNAR signal transduction^14^, was greatly diminished in *ISG15*-depleted cells upon EMCV infection as compared to infected cells that were transfected with the nontargeting control siRNA or MDA5-specific siRNA (**Extended Data Fig. 5b,d**), which is consistent with the reported role of ISG15 in preventing the degradation of USP18^15^. Together, these data suggested that in experimental settings of *ISG15*-gene targeting (*i.e. ISG15* gene silencing or KO) the antiviral effect of MDA5 ISGylation is masked by aberrant ISG upregulation due to the ablation of ISG15’s inhibitory effect on IFNAR-signal transduction.

**Figure 5.**
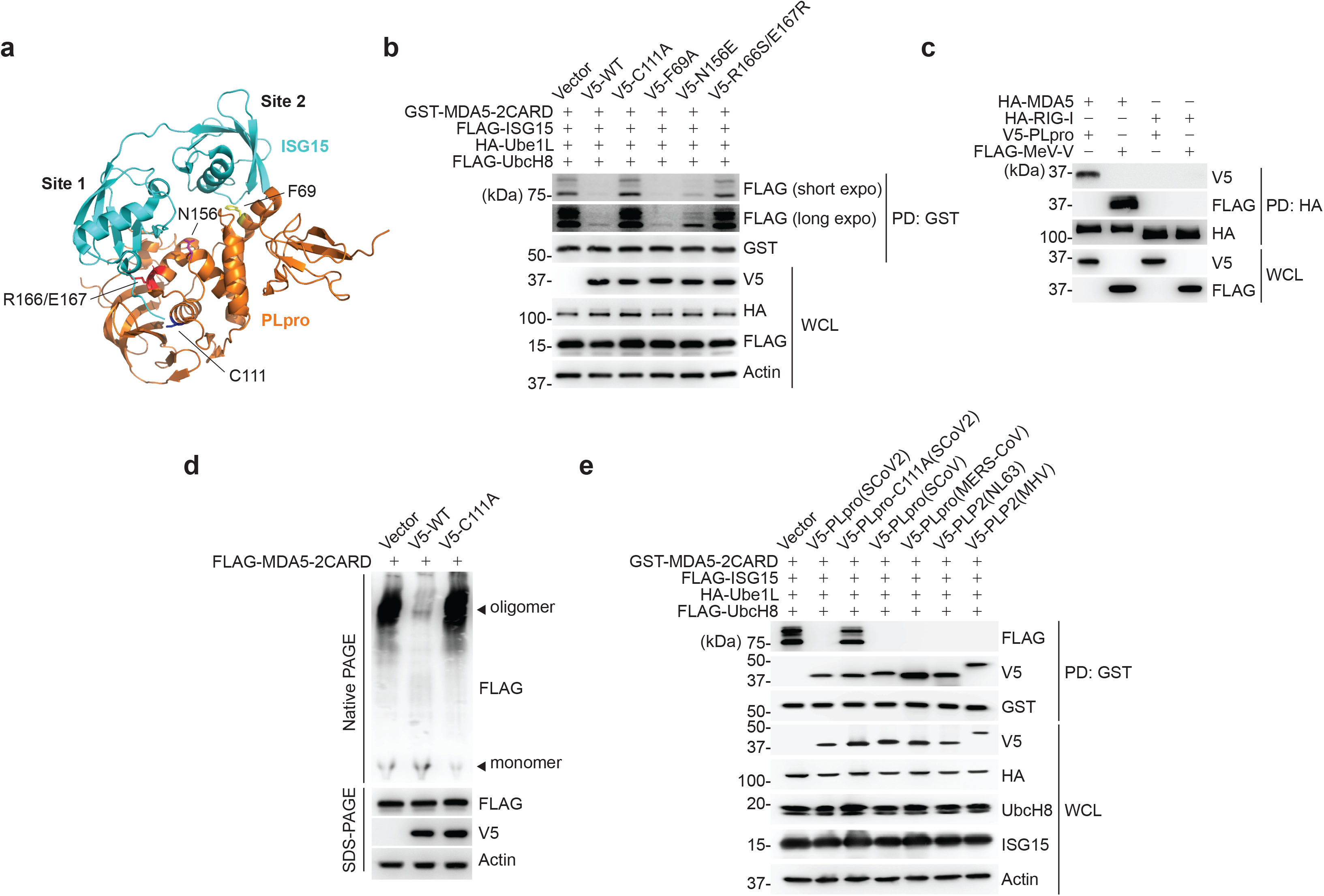
SCoV2 PLpro binds to and de-ISGylates MDA5-2CARD. **(a)** Ribbon representation of the crystal structure of the SCoV2 PLpro: ISG15 complex (PDB: 6YVA). Key residues that mediate ‘site 1’ interaction (N156 and R166/E167) or ‘site 2’ interaction (F69) in PLpro, as well as its catalytically-active site (C111), are indicated. **(b)** ISGylation of GST-MDA5-2CARD in HEK293T cells that were co-transfected for 20 h with vector or V5-tagged SCoV2 PLpro WT or mutants, along with FLAG-ISG15, HA-Ube1L, and FLAG-UbcH8, determined by GST-PD and IB with anti-FLAG and anti-GST. WCLs were probed by IB with anti-V5, anti-HA, anti-FLAG, and anti-Actin. **(c)** Binding of HA-tagged MDA5 or RIG-I to V5-SCoV2-PLpro or FLAG-MeV-V (control) in transiently transfected HEK293T cells, determined by HA-PD and IB with anti-V5 or anti-FLAG, and anti-HA. WCLs were probed by IB with anti-V5 and anti-FLAG. **(d)** Oligomerization of FLAG-MDA5-2CARD in HEK293T cells that were co-transfected with vector, or V5-SCoV2 PLpro WT or C111A for 24 h, assessed by Native PAGE and IB with anti-FLAG. WCLs were further analyzed by SDS-PAGE and probed by IB with anti-FLAG, anti-V5 and anti-Actin. **(e)** ISGylation of GST-MDA5-2CARD in HEK293T cells that also expressed FLAG-ISG15, HA-Ube1L and FLAG-UbcH8, and were co-transfected for 40 h with vector or the indicated V5-tagged coronaviral PLpro, determined by GST-PD and IB with anti-FLAG, anti-V5, and anti-GST. Data are representative of at least two independent experiments.

To test the effect of *ISG15* silencing on IFN-mediated virus restriction, we employed a virus protection assay that experimentally decouples MDA5 signaling in virus-infected cells from downstream IFNAR signaling in the same cells (**Fig. 4e**). Culture supernatants from mutEMCV-infected NHLF ‘donor’ cells that were either transfected with nontargeting control siRNA, or depleted of either *ISG15* or *MDA5* (positive control), were UV-inactivated and then transferred onto uninfected Vero ‘recipient’ cells. ‘Primed’ recipient cells were then infected with ZIKV to directly monitor the antiviral effect of MDA5-mediated IFN production by donor cells. Whereas the supernatants from control siRNA-transfected donor cells potently inhibited ZIKV replication, the supernatants from *ISG15* or *MDA5* knockdown cells minimally restricted virus replication (**Fig. 4f**). Consistent with these data, the culture supernatant from EMCV-infected HEK293 ‘donor’ cells that were transfected with WT MDA5 together with control siRNA led to greater protection of Vero ‘recipient’ cells from viral challenge than that from cells expressing WT MDA5 and depleted of *ISG15* (**Fig. 4g**). Collectively, these data demonstrate that ISGylation is important for MDA5-mediated restriction of a range of RNA viruses.

### SARS-CoV-2 PLpro targets MDA5 for de-ISGylation

Members of the *Coronaviridae* family, including SARS-CoV (SCoV), MERS-CoV, and the recently emerged SCoV2, encode a papain-like protease (PLpro) that, together with the main protease, mediates the cleavage of viral polyproteins^34^. In addition, PLpro has both deubiquitinating and de-ISGylating activities, which have been proposed to have immunomodulatory effects. A recent study showed that PLpro from SCoV2 modulates antiviral responses primarily via its de-ISGylase activity^20^; however, *bona fide* substrate(s) that are de-ISGylated by SCoV2 PLpro remain largely unknown. Since MDA5 is known to be a major sensor for detecting coronavirus infection^35, 36^, and because our data showed that ISGylation is essential for MDA5-mediated restriction of SCoV2 infection (**Fig. 4d**), we examined whether SCoV2 PLpro enzymatically removes the MDA5 CARD ISGylation to antagonize innate immunity. WT PLpro from SCoV2, but not its catalytically-inactive mutant C111A (PLpro-C111A)^20^, abolished MDA5-2CARD ISGylation (**Fig. 5a,b**). The PLpro N156E and R166S/E167R mutants, which are marginally and severely impaired in ISG15 binding at the ‘site 1’ interface^19, 37^, respectively, did slightly, or not, affect MDA5-2CARD ISGylation (**Fig. 5a,b**). In contrast, a SCoV2 PLpro mutant harboring the F69A mutation, which disrupts the ‘site 2’ interface that preferentially determines binding to ubiquitin, but not ISG15^19, 37^, diminished MDA5-2CARD ISGylation as potently as WT PLpro (**Fig. 5a,b**). SCoV2 PLpro, however, did not suppress RIG-I-2CARD ubiquitination; GST-RIG-I-2CARD was efficiently ubiquitinated in cells co-expressing WT PLpro or its catalytically-inactive mutant (C111A) (**Extended Data Fig. 6a**), which is in agreement with previous findings that indicated that SCoV2 PLpro has high specificity for cleaving K48-linked polyubiquitin, but not K63-ubiquitin linkages^19^.

**Figure 6.**
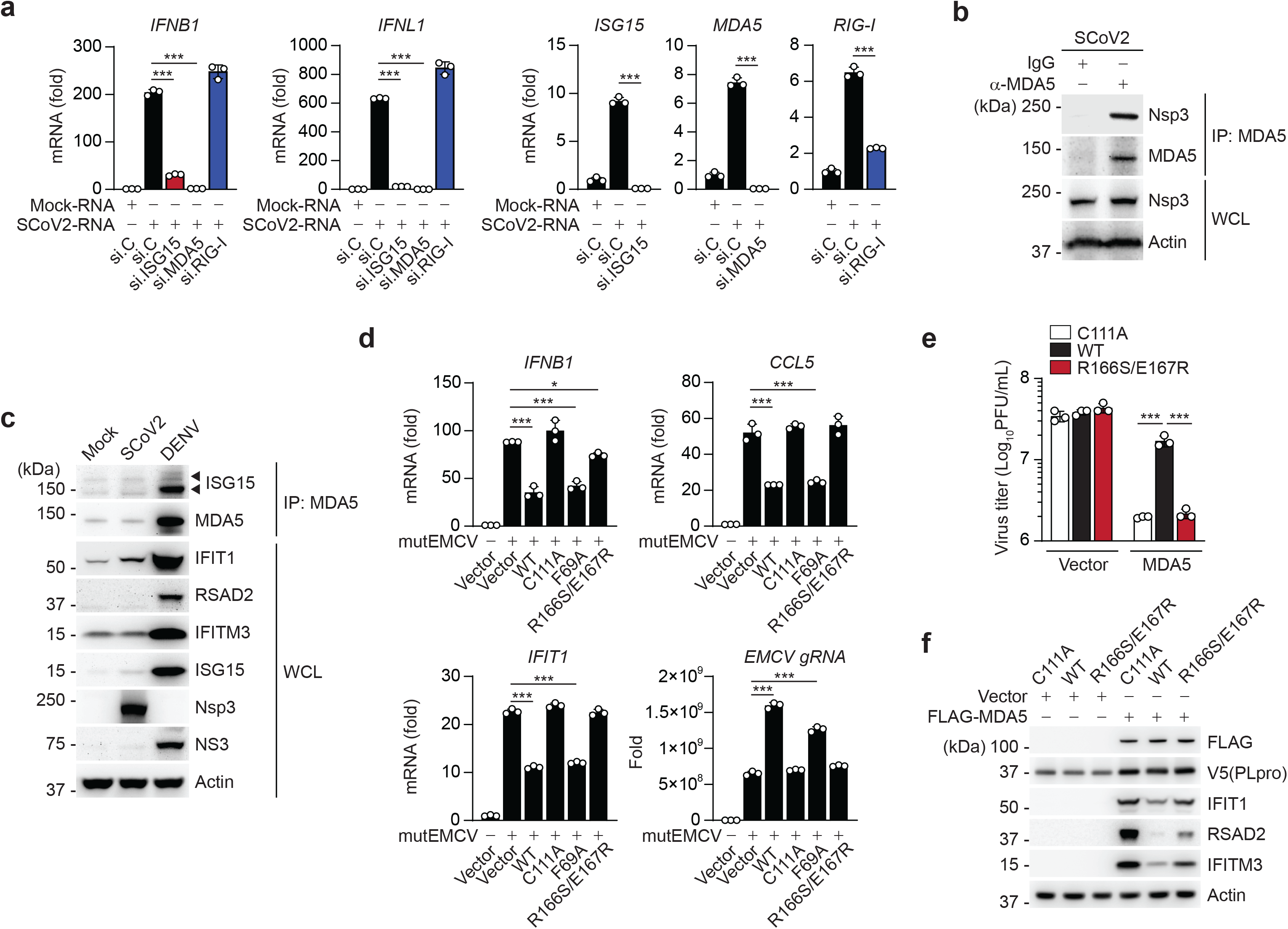
SCoV2 PLpro inhibits ISG15-mediated MDA5 signaling via its deISGylase activity. **(a)** qRT-PCR analysis of *IFNB1*, *IFNL1*, *ISG15*, *MDA5,* and *RIG-I* transcripts in NHLFs that were transfected with the indicated siRNAs for 40 h and then transfected with mock-RNA or SCoV2-RNA (0.4 μg/mL) for 24 h. **(b)** Binding of SCoV2 Nsp3 to endogenous MDA5 in A549-hACE2 cells that were infected with SCoV2 (MOI 0.5) for 24 h, determined by IP with anti-MDA5 (or an IgG isotype control) followed by IB with anti-PLpro and anti-MDA5. WCLs were probed by IB with anti-PLpro (Nsp3) and anti-Actin. **(c)** Endogenous MDA5 ISGylation in A549-hACE2 cells that were mock-infected, or infected with SCoV2 or DENV (MOI 0.1 for each) for 48 h, determined by immunoprecipitation (IP) with anti-MDA5 followed by IB with anti-ISG15 and anti-MDA5. Protein abundance of IFIT1, RSAD2, IFITM3, ISG15 and actin in the WCLs were probed by IB. Efficient virus replication was verified by immunoblotting WCLs with anti-PLpro (Nsp3) or anti-NS3 (DENV). **(d)** qRT-PCR analysis of *IFNB1*, *CCL5*, *IFIT1* transcript, and EMCV genomic RNA (gRNA) in HeLa cells that were transiently transfected for 24 h with vector, or V5-SCoV2 PLpro WT or mutants and then infected with mutEMCV (MOI 0.5) for 12 h. **(e)** EMCV titers in the supernatant of *RIG-I* KO HEK293 cells that were transiently transfected for 24 h with vector or FLAG-MDA5 along with V5-SCoV2 PLpro WT, C111A, or R166S/E167R and then infected with EMCV (MOI 0.001) for 16 h, determined by plaque assay. **(f)** Protein abundance of the indicated ISGs in the WCLs from the experiment in (e), determined by IB with the indicated antibodies. Data are representative of at least two independent experiments (mean ± s.d. of *n* = 3 biological replicates in a, d, e). \**p* < 0.05, \*\*\**p* < 0.001 (unpaired Student’s *t*-test).

In support of a direct activity of SCoV2 PLpro towards MDA5, we found that PLpro interacted specifically with MDA5, but not RIG-I, as did MeV-V that is known to bind MDA5 and therefore served as control^38^ (**Fig. 5c**). We found that low amounts of PLpro inhibited the signaling mediated by MDA5, but not by RIG-I, whereas higher amounts of PLpro suppressed antiviral signaling by both RLRs (**Extended Data Fig. 6b**). These results show that the MDA5 pathway is preferentially antagonized by PLpro, and also strengthen that MDA5 is a direct target of PLpro-mediated de-ISGylation. De-ISGylation of IRF3 likely accounts for the inhibitory effect that higher doses of PLpro have on the signaling by both sensors^39^.

To determine the consequence of MDA5 de-ISGylation by PLpro, we examined the effect of SCoV2 PLpro on MDA5-2CARD oligomerization. Ectopic expression of WT PLpro, similarly to *ISG15* depletion (**Fig. 3d**), efficiently blocked MDA5-2CARD oligomerization; in contrast, MDA5-2CARD efficiently oligomerized in cells co-transfected with empty vector or the SCoV2 PLpro C111A mutant (**Fig. 5d**), indicating that SCoV2 PLpro inhibits the ISGylation-dependent MDA5 oligomer formation via its enzymatic activity.

The PLpro enzymes of the related beta-coronaviruses, SCoV, MERS-CoV, and murine hepatitis virus (MHV), as well as of HCoV-NL63 (NL63) of the *Alphacoronavirus* genus, also efficiently reduced MDA5-2CARD ISGylation (**Fig. 5e**), suggesting that MDA5 antagonism by the de-ISGylase activity of PLpro may be widely conserved among the *Coronaviridae* family. In support of this, pull-down assay showed that the PLpro of SCoV, MERS-CoV, NL63 and MHV also bound to the MDA5-2CARD (**Fig. 5e**), indicating that MDA5 is a substrate of coronaviral PLpro de-ISGylating enzymes.

### SARS-CoV-2 PLpro antagonizes ISGylation-dependent MDA5 signaling

We next determined the relevance of ISG15-dependent MDA5 signaling for antiviral cytokine induction elicited by SCoV2. Since SCoV2 infection is known to minimally induce type I IFNs due to effective viral antagonisms^40^, we isolated total RNA from SCoV2-infected cells and then re-transfected it into cells to stimulate innate immune signaling. Transfection of total RNA from mock-infected cells served as a control. SCoV2-RNA, but not RNA from mock-infected cells, robustly triggered *IFNB1* and *IFNL1* transcript induction in NHLFs transfected with control siRNA. In contrast, siRNA-mediated silencing of *ISG15* or *MDA5* markedly diminished antiviral gene expression triggered by SCoV2-RNA (**Fig. 6a**). Notably, knockdown of endogenous RIG-I did not adversely affect the antiviral gene expression elicited by SCoV2-RNA, indicating that SCoV2 RNA-PAMP(s) are primarily sensed by the ISG15-MDA5-mediated signaling pathway (**Fig. 6a**). Next, in support of our finding of a SCoV2 PLpro-MDA5 2CARD interaction, we found that the SCoV2 non-structural protein 3 (Nsp3), within which PLpro lies, readily interacted with endogenous MDA5 during authentic SCoV2 infection (**Fig. 6b**). Endogenous MDA5 ISGylation and downstream ISG induction in SCoV2-infected cells were low (levels were similar to that in uninfected cells) as compared to those during DENV infection (**Fig. 6c**), supporting that PLpro effectively suppresses MDA5 ISGylation and innate signaling during live SCoV2 infection.

To provide evidence that SCoV2 PLpro antagonizes MDA5 signaling via its de-ISGylase activity, we examined the effect of WT and mutant PLpro on the activation of endogenous MDA5 during mutEMCV infection. Consistent with their effect on MDA5-2CARD ISGylation (**Fig. 5b**), ectopic expression of SCoV2 PLpro WT or the F69A mutant inhibited *IFNB1*, *CCL5* and *IFIT1* gene expression, whereas the de-ISGylase-deficient R166S/E167R mutant, similarly to the catalytically-inactive C111A mutant, did not affect the antiviral gene expression (**Fig. 6d**). In accord, the replication of mutEMCV was enhanced in cells expressing WT or F69A PLpro, but not in cells expressing C111A or R166S/E167R PLpro (**Fig. 6d**). Likewise, WT PLpro, but not the R166S/E167R or C111A mutant, blocked EMCV restriction by FLAG-MDA5 expression (**Fig. 6e**). The effect of the respective PLpro proteins on virus restriction correlated with the induction of ISG protein expression (*i.e*. IFIT1, RSAD2, and IFITM3) (**Fig. 6f**). Collectively, these results establish SCoV2 PLpro as an effective IFN antagonist that suppresses MDA5-mediated antiviral immunity via its de-ISGylase activity.

## DISCUSSION

ISG15, and in particular ISG15 conjugation, has long been known to confer antiviral activity to a multitude of viruses; however, only very few *bona fide* substrates (both viral and host-derived) have been identified^13^. On the other hand, ISG15 in its unconjugated form was shown to act provirally by negatively regulating USP18-mediated inhibition of IFNAR signal transduction^14, 15, 16^. Therefore, the physiological role of ISG15 in antiviral immunity has been elusive. This study shows that ISGylation of the viral RNA sensor MDA5 is crucial for its ability to elicit cytokine induction, demonstrating a key role of ISG15 in the IFN-mediated antiviral response. Whereas our study provided mechanistic insight into how ISGylation promotes antiviral innate immune responses, it is very likely that the sum of multiple ISGylation events (affecting both host and viral proteins) will ultimately determine the outcome of infection and pathogenesis, which may be context-dependent. Indeed, the effect of global ISG15 deficiency (*e.g.* using *ISG15* KO cells or mice) on virus replication has been extensively studied, which showed virus-specific effects^13^. Related to this, our work demonstrates an important framework of experimental design in which decoupling the role of ISG15 in MDA5 activation from that in downstream IFNAR signaling is essential to reveal ISG15’s antiviral function that acts through regulation of type I IFN production.

Whereas RIG-I activation is well known to require K63-linked ubiquitination^7^, MDA5 activation by PTMs is significantly less well understood. The MDA5 CARDs have been shown to be subject to several PTMs including non-covalent K63-linked polyubiquitin, SUMOylation, and K48-linked ubiquitination^5^. It will be important to investigate how specific PTMs are temporally regulated, or whether they have cell-type-specific roles. Along these lines, some PTMs may ‘cross-talk’ with each other. For example, whereas ISGylation at K23 and K43 promotes MDA5 activation during viral infection, degradative K48-linked ubiquitination at these sites may destabilize the MDA5 protein after the virus has been cleared successfully. Furthermore, how SUMOylation, which precedes PP1-mediated CARD dephosphorylation, influences MDA5 ISGylation, warrants further investigation. Regardless, our findings indicate that ISGylation of MDA5 acts analogously to the K63-linked ubiquitination of RIG-I in driving CARD-dependent RLR-signal activation. Similar to K63-ubiquitination of the RIG-I CARDs, ISGylation is dependent on PP1-mediated CARD dephosphorylation and promotes MDA5 CARD oligomerization and higher-order MDA5 assemblies. Despite these functional commonalities, ubiquitin and ISG15 have very distinct characteristics. Most notably, while ubiquitin is abundant in both uninfected and infected cells, ISG15 expression is minimal under normal (uninfected) conditions but is profoundly upregulated in response to IFN. Accordingly, the ISGylation of the host proteome is strongly increased in response to viral infection/IFN stimulation; however, even at basal levels, ISG15 is conjugated to many host proteins, including MDA5 as our work showed^17^. Thus, in viral infections exclusively sensed by MDA5, the low basal ISGylation activity of the cell may be sufficient for immediate activation of MDA5, whose basal levels are also extremely low. During viral infections that are sensed by multiple PRRs, MDA5 ISGylation may be a “priming” mechanism by which IFN induction and the ensuing ISG15 upregulation by immediate innate sensors (*e.g.* RIG-I) primes MDA5 to enter a ‘kick-start’ mode. This concept would be consistent with the temporal role of RIG-I and MDA5 during certain viral infections (for example, flaviviruses such as WNV) where RIG-I acts early and MDA5 signals later^41^. Interestingly, unlike MDA5, RIG-I has been shown to be negatively regulated by ISG15 (both covalent and noncovalent ISG15 binding has been reported)^21, 22^. Therefore, it is conceivable that the differential regulation of RIG-I and MDA5 by ISG15 may represent a mechanism of ‘sensor switching’ where MDA5 activation is promoted when ISG15 levels increase while, at the same time, RIG-I activity is being dampened.

The K63-linked ubiquitination of RIG-I is antagonized by several viral pathogens using a variety of mechanisms^42^. We identified that SCoV2 PLpro antagonizes MDA5 ISGylation (but not RIG-I CARD K63-ubiquitination) via its enzymatic activity after binding to the sensor. Our data also suggest that this immune evasion mechanism is likely conserved among several coronaviruses, which needs to be further investigated in the context of authentic infection. Interestingly, recent cryo-EM analyses revealed that the coronaviral Nsp3 protein is part of a molecular pore complex that spans ER-derived double-membrane vesicles and exports newly-synthesized viral RNA^43^. These results, combined with our findings identifying the MDA5-Nsp3(PLpro) interaction, support a model in which MDA5 may position itself in close proximity to the site of viral RNA export to facilitate PAMP detection; however, the PLpro domain of Nsp3 (which is on the cytoplasmic side) disarms MDA5 signaling function through direct de-ISGylation. Instead of direct de-ISGylation, some viruses may indirectly regulate MDA5 ISGylation, such as through manipulation of MDA5 S88 phosphorylation, as seen for the MeV V protein. Future studies should investigate the mechanistic details of viral evasion of ISG15-dependent MDA5 activation and, more broadly, the ISGylome manipulated by different viral pathogens that determines pathogenesis.

Taken together, our study uncovers a prominent role for ISGylation in activating MDA5-mediated immunity as well as its inhibition by SARS-CoV-2, unveiling a potential molecular target for the design of therapeutics against COVID-19.

## ACKNOWLEDGEMENTS

We greatly thank Deborah Lenschow (Washington University in St. Louis), Elmar Schiebel (University of Heidelberg), Ellen Cahir-McFarland (Biogen), Jan Rehwinkel (University of Oxford), Frank J.M. van Kuppeveld (Utrecht University), Karl-Klaus Conzelmann (LMU, Munich), Stephen Goodbourn (University of London), Susan C. Baker (Loyola University Chicago), Benjamin R. tenOever (Icahn School of Medicine at Mount Sinai), Adolfo García-Sastre (Icahn School of Medicine at Mount Sinai), and Jae U. Jung (Cleveland Clinic Lerner Research Institute) for providing reagents. We are also grateful to Sara Tavakoli and Jessica Poole for their help in the BSL-3 facility at the Cleveland Clinic Florida Research and Innovation Center. This study was supported in part by the US National Institutes of Health grants R01 AI087846 and R01 AI127774 (to M.U.G.).

## AUTHOR CONTRIBUTIONS

G.L., J-H.L., Z.M.P., M.U.G. designed the experiments; G.L., J-H.L., Z.M.P., D.A., M.v.G., W.R., J.J.C., M.E.D-G., E.W., and C.C. performed the experiments; G.L., J-H.L., Z.M.P., D.A., M.v.G., W.R., and M.U.G. analyzed data; M.U.G conceived the study; G.L., J-H.L., Z.M.P. and M.U.G. wrote the manuscript with input from all authors.

## ONLINE METHODS

### Cell culture

HEK293T (human embryonic kidney cells), Vero (African green monkey kidney epithelial cells), BHK-21 (Baby hamster kidney), and *Aedes albopictus* clone C6/36 cells were purchased from ATCC. Human peripheral blood monocuclear cells (PBMCs) were isolated from unidentified healthy donor peripheral blood (HemaCare) and purified by Lymphoprep density gradient centrifugation (STEMCELL Technologies). The WT and isogenic *Isg15*−/− MEFs (mouse embryonic fibroblasts) were kindly provided by Deborah Lenschow (Washington University in St. Louis). SVGAs (human fetal glial astrocytes) were kindly provided by Ellen Cahir-McFarland (Biogen)^1^. SVGA *MDA5* KO cells were generated by CRISPR/Cas9-mediated genome editing using a guide RNA (5'-AACTGCCTGCATGTTCCCGG-3') targeting the exon 1 of *IFIH1/MDA5*. The *MDA5* KO and *RIG-I* KO HEK293 cells were a gift from Jan Rehwinkel (University of Oxford)^2^. The WT and isogenic *ISG15* KO HeLa cells were kindly provided by Elmar Schiebel (University of Heidelberg)^3^. *ISG15* KO HeLa cells stably expressing FLAG-ISG15 WT or FLAG-ISG15 AA (GG156/157AA) were generated by lentiviral transduction followed by selection with puromycin (2 μg/mL). HAP-1 WT and isogenic *ISG15* KO cells were purchased from Horizon Discovery. HEK293T-hACE2 and Vero-E6-hACE2 were a gift from Jae U. Jung (Cleveland Clinic). A549-hACE2 were kindly provided by Benjamin R. tenOever (Icahn School of Medicine at Mount Sinai)^4^. HEK293T, HEK293, HeLa, MEFs, NHLFs, Vero, A549-hACE2, and BHK-21 cells were maintained in Dulbecco’s Modified Eagle’s Medium (DMEM, Gibco) supplemented with 10% (v/v) fetal bovine serum (FBS, Gibco), 2 mM GlutaMAX (Gibco), 1 mM sodium pyruvate (Gibco), and 100 U/mL penicillin-streptomycin (Gibco). HEK293T-hACE2 and Vero-E6-hACE2 were maintained in DMEM containing 200 μg/mL hygromycin B and 2 μg/mL puromycin respectively. SVGA and HAP-1 cells were cultured in Eagle’s Minimum Essential Medium (MEM, Gibco) and Iscove’s Modified Dulbecco’s Medium (IMDM, Gibco), respectively, supplemented with 10% FBS and 100 U/mL penicillin-streptomycin. PBMCs were maintained in RPMI 1640 (Gibco) supplemented with 10% FBS and 100 U/mL penicillin-streptomycin. C6/36 cells were cultured in MEM with 10% FBS and 100 U/mL penicillin-streptomycin. Except for C6/36 cells that were maintained at 28°C, all cell cultures were maintained at 37°C in a humidified 5% CO2 atmosphere.

### Viruses

DENV (serotype 2, strain 16681) and ZIKV (strain BRA/Fortaleza/2015) were propagated in C6/36 and Vero cells, respectively^5, 6^. Encephalomyocarditis virus (EMCV, EMC strain) was purchased from ATCC and propagated in HEK293T cells^7^. mutEMCV (EMCV-Zn_C19A/C22A_), which carries two point mutations in the zinc domain of the L protein^8^, was kindly provided by Frank J.M. van Kuppeveld (Utrecht University) and was propagated in BHK-21 cells. Sendai virus (strain Cantell) was purchased from Charles River Laboratories. SCoV2 (strain 2019-nCoV/USA_WA1/2020) was kindly provided by Jae U. Jung (Cleveland Clinic Lerner Research Center) and was propagated in Vero E6-hACE2 cells. All work relating to SCoV2 live virus and SCoV2-RNA was conducted in the BSL-3 facility of the Cleveland Clinic Florida Research and Innovation Center in accordance with institutional biosafety committee (IBC) regulations.

### DNA constructs and transfection

The human MDA5 ORF containing an N-terminal FLAG tag was amplified from the pEF-Bos-FLAG-MDA5^7^ and subcloned into pcDNA3.1/Myc-His B between *Xho*I and *Age*I. Site-directed mutagenesis on pcDNA3.1-FLAG-MDA5 (K23R/K43R, S88A, S88E, I841R/E842R, D848A/F849A, and G74A/W75A) was introduced by overlapping PCR. HA-MDA5 was cloned into pcDNA3.1(+) between *Kpn*I and *Xho*I. GST-MDA5-2CARD (in pEBG vector) and its S88A, S88D, S88E derivatives were described previously^7^. The single (K23R, K43R, K68R, K128R, K137R, K169R, K174R, and K235R) and double (K23R/K43R) mutations of MDA5-2CARD (aa 1-295) were introduced by site-directed mutagenesis into GST-MDA5-2CARD. Additionally, MDA5-2CARD and its K23R/K43R mutant were subcloned into pcDNA3.1(-) harboring an N-terminal 3×FLAG tag between *Nhe*I and *Not*I. pCR3-FLAG-MV-V (strain Schwarz) was a gift from Karl-Klaus Conzelmann (LMU, Munich). pEF-Bos-FLAG-NiV-V, pCAGGS-HA-MeV-V, and pCAGGS-HA-MeV-VΔtail were described previously^9^. Myc-tagged PIV2-V, MenV-V, MPRV-V, and HeV-V constructs were kindly provided by Stephen Goodbourn (University of London). FLAG-tagged PIV5-V, PIV2-V, MenV-V, MPRV-V, and HeV-V were subcloned into pEF-Bos containing an N-terminal FLAG tag between *Not*I and *Sal*I. pEF-Bos-FLAG-MuV-V was a gift from Curt Horvath (Addgene #44908^10^). pCAGGS-V5-hISG15 was a gift from Adolfo García-Sastre (Icahn School of Medicine at Mount Sinai)^11^. pCAGGS-HA-Ube1L and pFLAG-CMV2-UbcH8 were kindly provided by Jae U. Jung (University of Southern California). pcDNA3.1-Myc-UBE2I was cloned by ligating a synthetic UBE2I ORF into pcDNA3.1/Myc-His B between *Hind*III and *Not*I. FLAG-SUMO1 was obtained from Florian Full (University of Erlangen-Nuremberg, Germany). V5-tagged SARS-CoV-PLpro, MERS-CoV-PLpro, NL63-PLpro, MHV-PLP2 in pcDNA3.1-V5/His-B were kindly provided by Susan C. Baker (Loyola University of Chicago). The SARS-CoV-2 PLpro ORF (aa. 746-1060) was amplified from pDONR207 SARS-CoV-2 NSP3 (a gift from Fritz Roth; Addgene # 141257^12^) and subcloned into pcDNA3.1-V5. The C111A, F69A, N156E, R166S/E167R mutations of SARS-CoV-2-PLpro were introduced by site-directed mutagenesis. The correct sequence of all constructs was confirmed by DNA sequencing. Transient DNA transfections were performed using linear polyethylenimine [1 mg/mL solution in 10 mM Tris-HCl (pH 6.8); Polysciences], Lipofectamine 2000 (Invitrogen), Lipofectamine LTX with Plus Reagent (Invitrogen), *Trans*IT-HeLaMONSTER (Mirus), or *Trans*IT-X2 Transfection Reagent (Mirus) as per the manufacturers’ instructions.

### Antibodies and other reagents

Primary antibodies used in this study include anti-GST (1:5,000; Sigma-Aldrich), anti-V5 (1:5,000, R960-25; Novex), anti-FLAG (M2, 1:2,000; Sigma-Aldrich), anti-HA (1:3,000, HA-7; Sigma-Aldrich), anti-Phospho-IRF-3 (Ser396) (1:1,000, D6O1M; CST), anti-IRF3 (1:1,000, D6I4C; CST), anti-Phospho-STAT1 (Tyr701) (1:1,000, 58D6; CST), anti-IFIT1 (1:1,000, PA3-848; Invitrogen and 1:1,000, D2X9Z; CST), anti-IFIT2 (1:1,000; Proteintech), anti-ISG15 (1:500, F-9; Santa Cruz), anti-MAVS (1:1,000; CST), anti-RIG-I (1:2,000, Alme-1; Adipogen), anti-MDA5 (1:1,000, D74E4; CST), anti-Phospho-MDA5 (Ser88)^7^, anti-PP1α (1:2,000; Bethyl laboratories), anti-PP1γ (1:2,000; Bethyl laboratories), anti-USP18 (1:1000, D4E7; CST), anti-RSAD2 (1:1,000, D5T2X; CST), anti-PKR (1:1,000, D7F7; CST), anti-MX1 (1:1,000, D3W7I; CST), anti-IFITM3 (1:1,000, D8E8G; CST), anti-ISG20 (1:1,000, PA5-30073; Invitrogen), anti-ubiquitin (1:1,000, P4D1; Santa Cruz), anti-NS3^6^, anti-PLpro (Nsp3) (1:1,000, GTX135589; GeneTex), anti-β-tubulin (1:1,000; CST), and anti-β-Actin (1:1,000, C4; Santa Cruz). Monoclonal anti-MDA5 antibody was purified from mouse hybridoma cell lines kindly provided by Jan Rehwinkel (University of Oxford)^2^. Monoclonal anti-IFNAR2 neutralizing antibody (1:250, MMHAR-2) was obtained from PBL Assay Science. Monoclonal anti-flavivirus E antibody (4G2) was purified from the mouse hybridoma cell line D1-4G2-4-15 (ATCC). Anti-mouse and anti-rabbit HRP-conjugated secondary antibodies (1:2,000) were purchased from CST. Anti-FLAG M2 magnetic beads (Sigma-Aldrich), anti-FLAG agarose beads (Sigma-Aldrich), Glutathione Sepharose 4B resin (GE Healthcare), and Protein G Dynabeads (Invitrogen) were used for protein immunoprecipitation. Protease and phosphatase inhibitors were obtained from Sigma-Aldrich. Poly(I:C) (HMW)/LyoVec and Poly(I:C) (HMW) Biotin were obtained from Invivogen. Human IFN-β was purchased from PBL Biomedical Laboratories.

### Mass spectrometry

Large-scale GST-pulldown and mass spectrometry (MS) analysis were performed as previously described^13, 14^. Briefly, HEK293T cells were transfected with GST or GST-MDA5-2CARD, and the cells were collected at 48 h post-transfection and lysed in Nonidet P-40 (NP-40) buffer [50 mM HEPES (pH 7.4), 150 mM NaCl, 1% (v/v) NP-40, 1 mM EDTA, and 1× protease inhibitor cocktail (Sigma)]. Cell lysates were cleared by centrifugation at 20,000 ×*g* at 4°C for 20 min, and cleared supernatants were subjected to GST-pulldown using glutathione Sepharose 4B beads (GE Healthcare) at 4°C for 4 h. The beads were extensively washed with NP-40 buffer and proteins eluted by heating in 5× Laemmli SDS sample buffer at 95°C for 5 min. Eluted proteins were resolved on a NuPAGE 4-12% Bis-Tris gel (Invitrogen) and then stained at room temperature using the SilverQuest Silver Staining Kit (Invitrogen). The bands that were specifically present in the GST-MDA5-2CARD sample, but not the GST control sample, were excised and analyzed by LC-MS/MS (Taplin Mass Spectrometry Facility, Harvard University).

### Immunoprecipitation and immunoblotting

Cells were transfected with FLAG-MDA5, GST-MDA5-2CARD, or FLAG-MDA5-2CARD in the absence or presence of ISGylation machinery components (*i.e.* HA-Ube1L, FLAG-UbcH8, and V5-ISG15) as indicated. Forty-eight hours later, cells were lysed in NP-40 buffer and cleared by centrifugation at 20,000 ×*g* at 4°C for 20 min. Cell lysates were then subjected to GST or FLAG pulldown using glutathione magnetic agarose beads (Pierce) and anti-FLAG M2 magnetic beads (Millipore) at 4°C for 4 h or 16 h respectively. The beads were extensively washed with NP-40 buffer and proteins eluted by heating in 1× Laemmli SDS sample buffer at 95°C for 5 min or by competition with FLAG peptide (Millipore) 4°C for 4 h. For endogenous MDA5 immunoprecipitation, NHLFs were stimulated with poly(I:C) (HMW)/LyoVec (0.1 μg/mL) or infected with DENV or ZIKV at the indicated MOI for 40 h. Cell lysates were precleared with Protein G Dynabeads (Invitrogen) at 4°C for 2 h and then incubated with Protein G Dynabeads conjugated with the anti-MDA5 antibody or an IgG1 isotype control (G3A1; CST) at 4°C for 4 h. The beads were washed four times with RIPA buffer [20 mM Tris-HCl (pH 8.0), 150 mM NaCl, 1% (v/v) NP-40, 1% (w/v) deoxycholic acid, 0.01% (w/v) SDS] and protein eluted in 1× Laemmli SDS sample buffer. Protein samples were resolved on Bis-Tris SDS-PAGE gels, transferred onto polyvinylidene difluoride (PVDF) membranes (Bio-Rad), and visualized using the SuperSignal West Pico PLUS or Femto chemiluminescence reagents (Thermo Scientific) on an ImageQuant LAS 4000 Chemiluminescent Image Analyzer (General Electric) as previously described^6^.

### Enzyme-linked immunosorbent assay (ELISA)

Human or mouse IFN-β in the culture supernatants of NHLFs, HeLa, and MEFs was determined by ELISA using the VeriKine Human Interferon Beta ELISA Kit or VeriKine Mouse Interferon Beta ELISA Kit (PBL Assay Science) as previously described^7^.

### siRNA- and shRNA-mediated knockdown

Transient knockdown in NHLFs, HeLa, HAP-1, HEK293T, and HEK293 cells was performed using non-targeting or gene-specific siGENOME SMARTpool siRNAs (Dharmacon). These are the Non-Targeting siRNA Pool #2 (D-001206-14), *IFIH1* (M-013041-00), *DDX58* (M-012511-01), *PPP1CA* (M-008927-01), *PPP1CC* (M-006827-00) and *ISG15* (D-004235-17 and D-004235-18). Transfection of siRNAs was performed using the Lipofectamine RNAiMAX Transfection Reagent (Invitrogen) as per the manufacturer’s instructions. Scrambled shRNA control lentiviral particles and shRNA lentiviral particles targeting *ISG15* (TL319471V) or *IFIH1* (TL303992V) were purchased from OriGene. Lentiviral transduction of human PBMCs (1 ×10^5^ cells; MOI 8) was performed in the presence of 8 μg/mL polybrene (Santa Cruz). Knockdown efficiency was determined by qRT-PCR or immunoblotting as indicated.

### Quantitative real-time PCR (qRT-PCR)

Total RNA was purified using the E.Z.N.A. HP Total RNA Kit (Omega Bio-tek) as per the manufacturer’s instructions. One-step qRT-PCR was performed using the SuperScript III Platinum One-Step qRT-PCR Kit (Invitrogen) and predesigned PrimeTime qPCR Probe Assays (IDT) on a 7500 Fast Real-Time PCR System (Applied Biosystems). Relative mRNA expression was normalized to the levels of *GAPDH* and expressed relative to the values for control cells using the ΔΔCt method.

### Luciferase reporter assay

IFN-β reporter assay was performed as previously described^15^. Briefly, HEK293T or *MDA5* KO HEK293 cells were transfected with IFN-β luciferase reporter construct and β-galactosidase (β-gal) expressing pGK-β-gal, along with GST-MDA5-2CARD (WT or mutants) or FLAG-MDA5 (WT or mutants). At the indicated time points after transfection, luciferase and β-gal activities were determined using respectively the Luciferase Assay System (Promega) and β-Galactosidase Enzyme Assay System (Promega) on a Synergy HT microplate reader (BioTek). Luciferase activity was normalized to β-gal values, and fold induction was calculated relative to vector-transfected samples, set to 1.

### Cytosol-mitochondria fractionation assay

The cytosol-mitochondria fractionation assay was performed using a Mitochondria/Cytosol Fractionation Kit (Millipore) as previously described^5, 6^. Briefly, NHLFs were transfected for 24 h with either non-targeting control siRNA or ISG15-specific siRNA and then transfected with EMCV-RNA or RABV_Le_ for 16 h. Cells were homogenized in an isotonic buffer using a Dounce homogenizer and the lysates were centrifuged at 600 ×*g* to pellet the nuclei and unbroken cells. The supernatant was further centrifuged at 10,000 ×*g* at 4°C for 30 min to separate the cytosolic (supernatant) and mitochondrial (pellet) fractions. The protein concentration of both fractions was determined by a bicinchoninic acid (BCA) assay (Pierce), and equal amounts of proteins were analyzed by immunoblotting. Anti-β-tubulin and anti-MAVS immunoblotting served as markers for the cytosolic and mitochondrial fractions respectively.

### *In vitro* RNA-binding assay

WT and *Isg15*−/− MEFs were stimulated with IFN-β (1,000 U/mL) for 24 h. Cells were lysed in a buffer containing 50 mM HEPES (pH 7.4), 200 mM NaCl, 1% (v/v) NP-40, 1 mM EDTA, and 1× protease inhibitor cocktail (Sigma). NeutrAvidin agarose beads (Pierce) were conjugated with the biotinylated HMW-Poly(I:C) at 4°C for 4 h. Cell lysates were incubated with the conjugated beads at 4°C for 16 h. The beads were washed three times with lysis buffer and then boiled at 95°C in 1× Laemmli SDS sample buffer to elute the proteins. Precipitated proteins were resolved on Bis-Tris SDS-PAGE gels and analyzed by IB with anti-MDA5. Equal input MDA5 protein amounts were confirmed by IB with anti-MDA5.

### Native PAGE

Native PAGE for analyzing endogenous IRF3 dimerization was performed as previously described^16^. For measuring MDA5 oligomerization, HEK293T or HEK293 *MDA5* KO cells were transfected with WT or mutant FLAG-MDA5-2CARD or FLAG-MDA5 as indicated. Twenty-four hours later, cells were lysed in 1× NativePAGE sample buffer (Invitrogen) containing 1% (v/v) NP-40 on ice for 30 min and then lysates were cleared by centrifugation at 16,000 ×*g* at 4°C for 10 min. Cleared lysates were resolved on a 3-12% Bis-Tris NativePAGE gel (Invitrogen) as per the manufacturer's instructions and analyzed by immunoblotting with the indicated antibodies.

### Semi-denaturating detergent agarose gel electrophoresis (SDD-AGE)

MDA5 oligomerization in MEFs transfected with EMCV-RNA, or in HEK293 *MDA5* KO cells reconstituted with WT or mutant FLAG-MDA5, was determined by SDD-AGE as previously described with modifications^17^. Briefly, cells were lysed in a buffer containing 50 mM HEPES (pH 7.4), 150 mM NaCl, 0.5% (v/v) NP-40, 10% (v/v) glycerol, and 1× protease inhibitor cocktail (Sigma) at 4°C for 20 min. Cell lysates were cleared by centrifugation at 16,000 ×*g* at 4°C for 10 min and then incubated on ice for 1 h. Cell lysates were subsequently incubated in 1× SDD-AGE buffer (0.5× TBE, 10% (v/v) glycerol, and 2% (w*/*v) SDS) for 15 min at room temperature and resolved on a vertical 1.5% agarose gel containing 1× TBE and 0.1% (w*/*v) SDS at 80 V for 90 min at 4°C. Proteins were transferred onto a PVDF membrane and analyzed by immunoblotting with the indicated antibodies.

### Viral RNA purification

EMCV-RNA was produced as previously described^7^. Briefly, Vero cells were infected with EMCV (MOI 0.1) for 16 h, and total RNA was isolated using the Direct-zol RNA extraction kit (Zymo Research) as per the manufacturer’s instructions. Mock-RNA and SCoV2-RNA were produced by isolating total RNA from uninfected or SCoV2-infected (MOI 1 for 24 h) Vero-hACE2 cells. RABV_Le_ was generated by *in vitro* transcription using the MEGAshortscript T7 Transcription Kit (Invitrogen) as previously described^18^.

### Virus infection and titration

All viral infections were performed by inoculating cells with the virus inoculum diluted in MEM or DMEM containing 2% FBS. After 1-2 h, the virus inoculum was removed and replaced with the complete growth medium (MEM or DMEM containing 10% FBS) and cells were further incubated for the indicated times. EMCV titration was performed either on Vero cells using the median tissue culture infectious dose (TCID50) methodology as previously described^19^, or on BHK-21 cells by the standard plaque assay. The titers of ZIKV were determined by plaque assay on Vero cells as previously described^6^. Titration of SCoV2 was performed on Vero-hACE2 cells by plaque assay.

### Flow cytometry

To quantify the percentage of DENV-infected cells, reconstituted HEK293 *MDA5* KO cells were washed with PBS (Gibco) and fixed with 4% (v/v) formaldehyde in PBS at room temperature for 30 min. Cells were subsequently permeabilized with 1× BD Perm/Wash buffer (BD Biosciences) for 15 min and incubated with an anti-flavivirus E antibody (4G2; 1:100 in 1× BD Perm/Wash buffer) at 4°C for 30 min. Cells were further washed three times with 1× BD Perm/Wash buffer and incubated with a goat anti-mouse Alexa Fluor 488-conjugated secondary antibody (#A10667, 1:500 in 1× BD Perm/Wash buffer; Invitrogen) at 4°C for 30 min in the dark. After washing three times with 1× BD Perm/Wash buffer, cells were analyzed on a FACSCalibur flow cytometer (BD Biosciences). Data analysis was performed using the FlowJo software.

### Virus protection assay

The culture supernatants from mutant or WT EMCV-infected NHLFs or *RIG-I* KO HEK293 cells were UV-inactivated in a biosafety cabinet under a UV-C lamp (30W) at a dose of 5,000 μJ/cm^2^ for 15 min. Complete inactivation of EMCV was confirmed by plaque assay on BHK-21 cells. The inactivated supernatants were then transferred onto fresh Vero cells for 24 h, and the primed Vero cells were subsequently infected with ZIKV (MOI 0.002 to 2) for 72 h, or with EMCV (MOI 0.001 to 0.1) for 40 h. ZIKV-positive cells were determined by immunostaining with anti-flavivirus E antibody (4G2) and visualized using the KPL TrueBlue peroxidase substrate (SeraCare). EMCV-induced cytopathic effect was visualized by Coomassie blue staining.

### Statistical analysis

Unpaired Student’s *t* test was used to compare differences between two experimental groups in all cases. Significant differences are denoted by \**p* < 0.05, \*\**p* < 0.01, or \*\*\**p* < 0.001. Pre-specified effect sizes were not assumed, and in general, three biological replicates (*n*) for each condition were used.

## DATA AVAILABILITY

The data that support the findings of this study are available from the corresponding author upon request.

