## Supplemental Figures 1-6 for "ISG15-dependent Activation of the RNA Sensor MDA5 and its Antagonism by the SARS-CoV-2 papain-like protease"

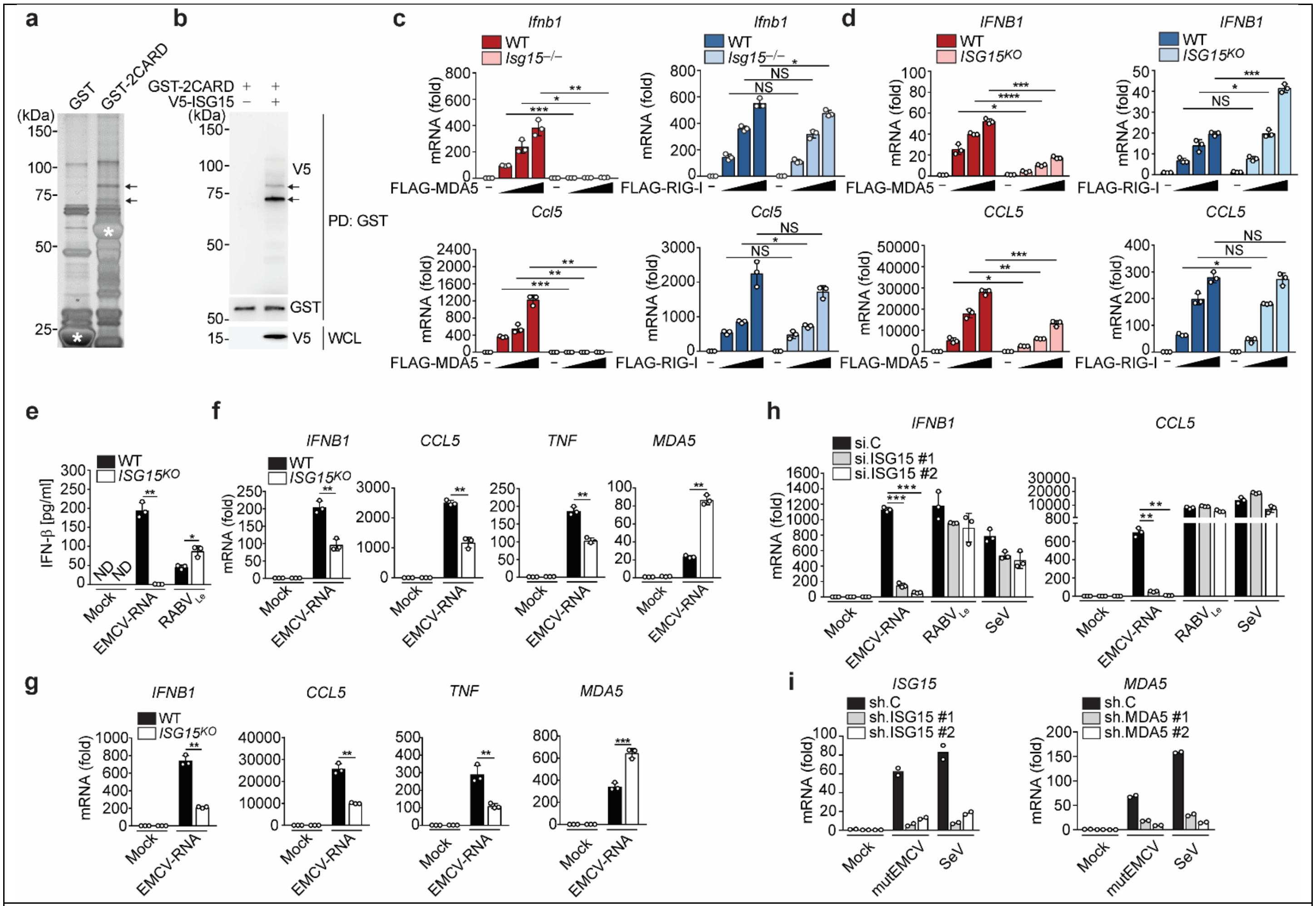


**Extended Data Fig. 1**

**ISG15 is required for MDA5, but not RIG-I, mediated signal transduction.**

**(a)** Silver-stained affinity-purified GST and GST-MDA5-2CARD from transiently transfected HEK293T cells. Asterisks denote the GST and GST-MDA5-2CARD (aa 1-295) proteins. Arrows indicate the bands that identified ISG15 by MS analysis. **(b)** GST-MDA5-2CARD ISGylation in transiently transfected HEK293T cells with or without co-expressed V5-ISG15, determined by GST pulldown (PD) and immunoblot (IB) with anti-GST and anti-V5. Whole cell lysates (WCLs) were probed by IB with anti-V5. **(c, d)** qRT-PCR analysis of *IFNB1* and *CCL5* transcripts in WT and *Isg15*$^{-/-}$ MEFs (c) or WT and *ISG15* KO HeLa cells (d) that were transfected with empty vector or increasing amounts of FLAG-MDA5 or FLAG-RIG-I for 40 h. **(e)** ELISA of IFN-β in the supernatants of WT or *ISG15* KO HeLa cells that were mock-transfected or transfected with EMCV-RNA (0.4 μg/mL) or $RABV_{Le}$ (1 pmol/mL) for 24 h. **(f)** qRT-PCR analysis of *IFNB1*, *CCL5*, *TNF*, and *MDA5* mRNA in WT and *ISG15* KO HeLa cells that were mock-transfected or transfected with EMCV-RNA (0.4 μg/mL) for 24 h. **(g)** qRT-PCR analysis of *IFNB1*, *CCL5*, *TNF*, and *MDA5* mRNA in WT and *ISG15* KO HAP-1 cells that were stimulated as in (f). **(h)** qPCR analysis of *IFNB1* and *CCL5* mRNA in NHLFs that were transfected with the indicated siRNAs for 30 h and then mock-stimulated or transfected with EMCV-RNA (0.4 μg/mL) or $RABV_{Le}$ (1 pmol/mL), or infected with SeV (10 HAU/mL) for 16 h. **(i)** qRT-PCR analysis of *ISG15* and *MDA5* mRNA in PBMCs that were transduced for 40 h with the indicated shRNA lentiviral particles and then infected with mutEMCV (MOI 10) or SeV (200 HAU/mL) for 8 h Data are representative of at least two independent experiments (mean ± s.d. of $n = 3$ biological replicates in c, d, e, f, and g; mean of $n = 2$ biological replicates in i). $^{*}p < 0.05$, $^{**}p < 0.01$, $^{***}p < 0.001$ (unpaired Student's *t*-test). ND, not detected.

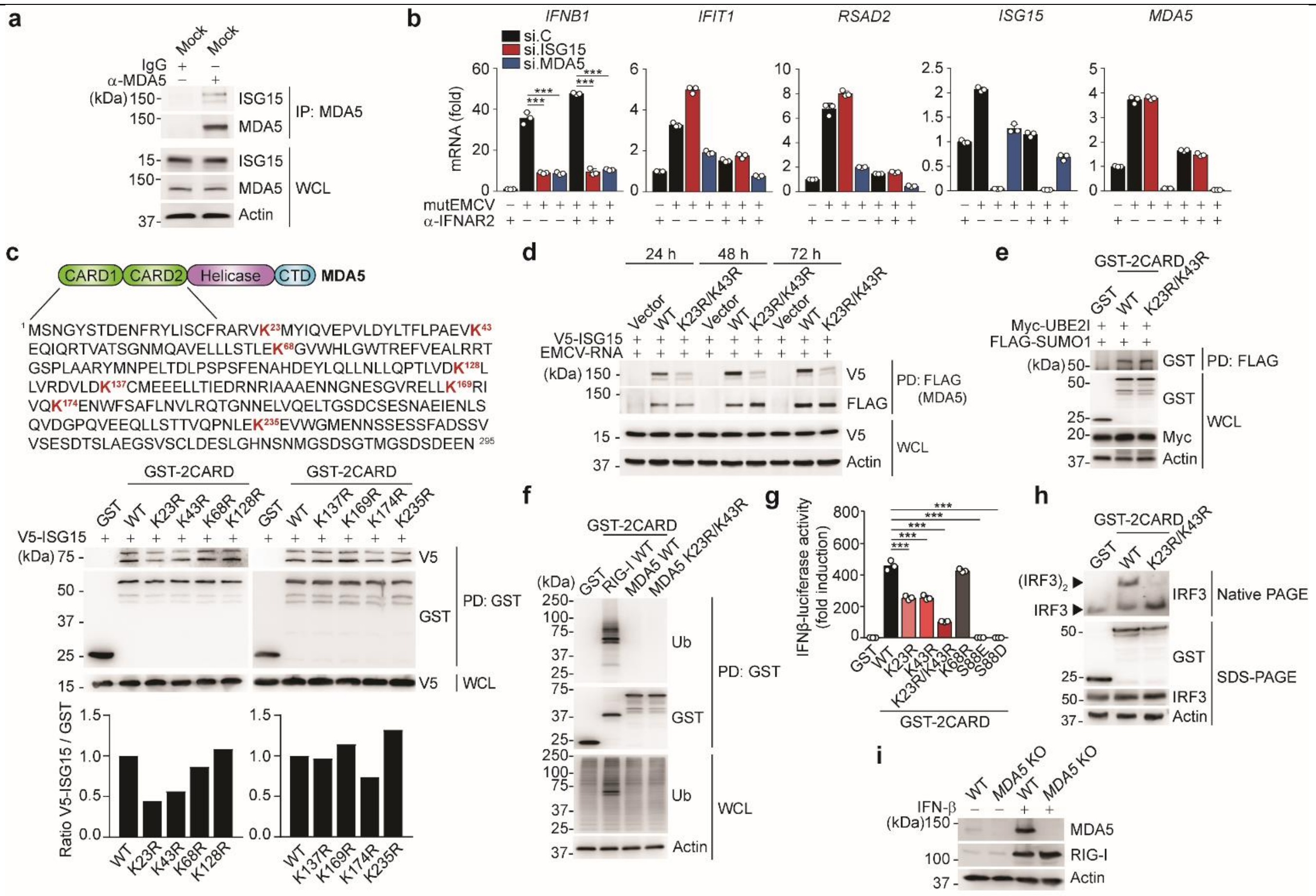

**Extended Data Fig. 2**

**ISGylation at K23 and K43 is essential for MDA5 activation.**

**(a)** ISGylation of endogenous MDA5 in uninfected NHLFs, determined by immunoprecipitation (IP) with anti-MDA5 (or IgG isotype control antibody) and IB with anti-ISG15 and anti-MDA5. WCLs were probed by IB with anti-ISG15, anti-MDA5, and anti-Actin. **(b)** qRT-PCR analysis of *IFNB1*, *IFIT1*, *RSAD2*, *ISG15* and *MDA5* mRNA in NHLFs that were transfected for 30 h with the indicated siRNAs and then infected with mutEMCV (MOI 0.005) for 16 h in the absence or presence of anti-IFNAR2 antibody (2 µg/mL). **(c)** Upper panels: MDA5 domain architecture and amino acid sequence of the CARDs. Lysine (K) residues within the CARDs are highlighted in red. CTD, C-terminal domain. Middle panels: ISGylation of GST-MDA5-2CARD WT and indicated K-to-R mutants in transiently transfected HEK293T cells that co-expressed V5-ISG15, determined by GST-PD and IB with anti-GST and anti-V5. WCLs were probed by IB with anti-V5. Lower panels: Densitometric analysis of ISGylation levels of the indicated K-to-R mutants, normalized to GST-PD levels. Data are presented as fold induction relative to the values for cells transfected with WT GST-MDA5-2CARD, set to 1. **(d)** ISGylation of FLAG-MDA5 WT and K23R/K43R in transiently transfected *MDA5* KO HEK293 cells that co-expressed V5-ISG15 and were stimulated with EMCV-RNA (0.4 µg/mL) for the indicated times, determined by FLAG-PD and IB with anti-FLAG and anti-V5. WCLs were probed by IB with anti-V5 and anti-Actin. **(e)** SUMOylation of GST-MDA5-2CARD WT and K23R/K43R in transiently transfected HEK293T cells that co-expressed Myc-UBE2I and FLAG-SUMO1, determined by FLAG-PD and IB with anti-GST. WCLs were probed by IB with anti-GST, anti-Myc, and anti-Actin. **(f)** Ubiquitination of GST-RIG-I-2CARD WT, GST-MDA5-2CARD WT and K23R/K43R in transiently transfected HEK293T cells, determined by GST-PD and IB with anti-Ub. WCLs were probed by IB with anti-GST, anti-Ub, and anti-Actin. **(g)** IFN-β-luciferase activity in HEK293T cells that were transfected for 40 h with GST, or GST-MDA5-2CARD (GST-2CARD) WT or mutants. Luciferase activity is presented as fold induction relative to the values for GST-transfected cells, set to 1. **(h)** Endogenous IRF3 dimerization in HEK293T cells that were transiently transfected with GST, GST-MDA5-2CARD WT or K23R/K43R for 24 h, determined by Native PAGE and IB with anti-IRF3. WCLs were additionally analyzed by SDS-PAGE and IB with anti-GST, anti-IRF3 and anti-Actin (loading control). **(i)** Validation of *MDA5* gene editing in *MDA5* KO SVGAs. Protein abundance of endogenous MDA5 in the WCLs of WT control or *MDA5* KO SVGA cells that were treated with IFN-β (1,000 U/mL) for 16 h, assessed by IB with anti-MDA5. WCLs were further probed by IB with anti-RIG-I and anti-Actin. Data are representative of at least two independent experiments (mean ± s.d. of $n$ = 3 biological replicates in b and g). ***$p < 0.001$ (unpaired Student's $t$-test).

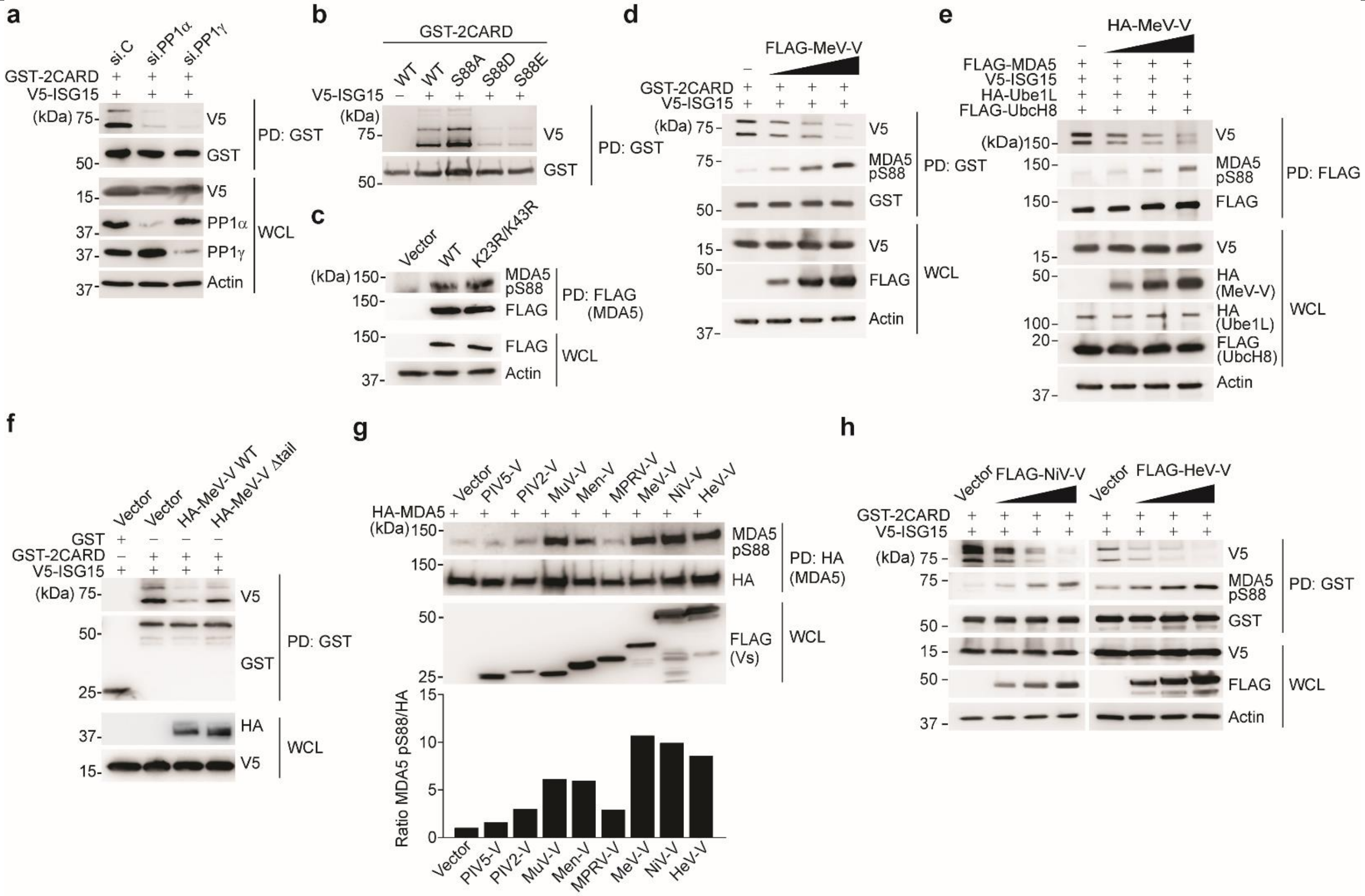


**Extended Data Fig. 3**

**Dephosphorylation of MDA5 induces ISGylation.**

**(a)** ISGylation of GST-MDA5-2CARD in HEK293T cells transfected with V5-ISG15 and the indicated siRNAs for 48 h, assessed by GST-PD and IB with anti-V5 and anti-GST. WCLs were probed by IB with anti-V5, anti-PP1α, anti-PP1γ, and anti-Actin. **(b)** ISGylation of GST-MDA5-2CARD WT or S88A, S88D and S88E mutants in transiently transfected HEK293T cells that also co-expressed V5-ISG15, determined by GST-PD and IB with anti-V5 and anti-GST forty hours after transfection. **(c)** Phosphorylation of FLAG-MDA5 WT and K23R/K43R in HEK293T cells, determined by FLAG-PD and IB with anti-pS88-MDA5 and anti-FLAG. WCLs were probed by IB with anti-FLAG and anti-Actin. **(d)** ISGylation and phosphorylation of GST-MDA5-2CARD in HEK293T cells transfected with V5-ISG15 and either empty vector or increasing amounts of FLAG-MeV-V for 24 h, determined by GST-PD and IB with anti-pS88-MDA5, anti-V5, and anti-GST. **(e)** ISGylation and phosphorylation of FLAG-tagged MDA5 in transiently transfected HEK293T cells that also co-expressed V5-ISG15, HA-Ube1L, and FLAG-UbcH8 as well as either empty vector or increasing amounts of HA-MeV-V for 24 h, determined by FLAG-PD and IB with anti-pS88-MDA5, anti-V5, and anti-FLAG. WCLs were probed by IB with the indicated antibodies. **(f)** ISGylation of GST-MDA5-2CARD in HEK293T cells transiently transfected with V5-ISG15 and either empty vector or HA-tagged MeV-V WT or Δtail for 48 h, determined by GST-PD and IB with anti-V5 and anti-GST. **(g)** Upper panel: Phosphorylation of HA-MDA5 in transiently transfected HEK293T cells that also co-expressed empty vector or the indicated FLAG-tagged paramyxoviral V protein, assessed by HA-PD and IB with anti-pS88-MDA5 and anti-HA. WCLs were probed by IB with anti-FLAG. Lower panel: Densitometric analysis of levels of MDA5 phosphorylation (pS88) that were normalized to HA-PD levels. Data are presented as fold induction relative to the values for cells transfected with HA-MDA5 and vector, set to 1. **(h)** ISGylation and phosphorylation of GST-MDA5-2CARD in transiently transfected HEK293T cells that also co-expressed V5-ISG15 and either empty vector or increasing amounts of FLAG-NiV-V or FLAG-HeV-V for 24 h, determined by GST PD and IB with anti-V5, anti-pS88-MDA5, and anti-GST. WCLs were probed by IB with anti-FLAG, anti-V5, and anti-Actin. Data are representative of at least two independent experiments.

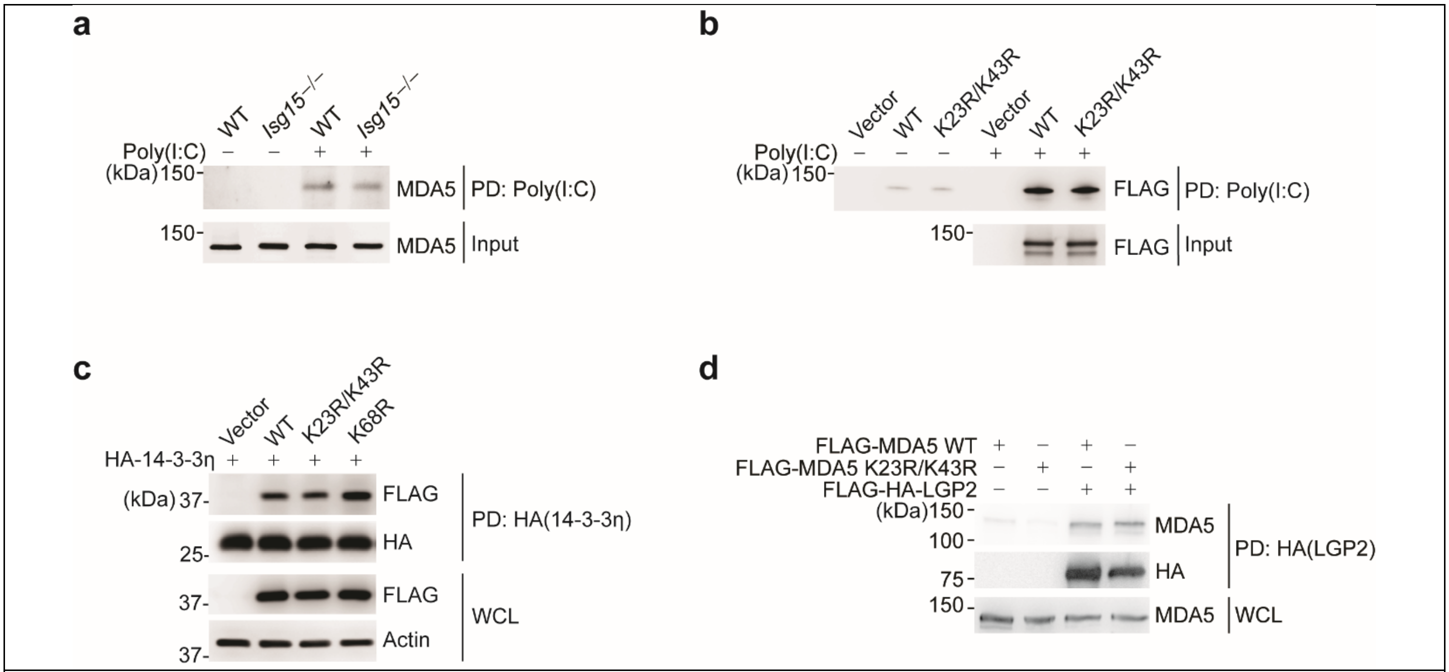


**Extended Data Fig. 4**

**CARD ISGylation does not affect the ability of MDA5 to bind RNA, 14-3-3η, or LGP2.**

**(a)** *In vitro* RNA-binding ability of endogenous MDA5 from WT or *Isg15*$^{-/-}$ MEFs that were stimulated with IFN-β (1,000 U/mL) for 24 h, assessed by biotin-HMW-poly(I:C)-PD and IB with anti-MDA5. Equal input MDA5 protein amounts were confirmed by IB with anti-MDA5. **(b)** *In vitro* RNA-binding ability of FLAG-MDA5 WT and K23R/K43R from transiently transfected HEK293T cells, assessed by biotin-HMW-poly(I:C)-PD and IB with anti-FLAG. Equal input FLAG-MDA5 protein amounts were confirmed by WCLs were probed by IB with anti-FLAG. **(c)** Binding of FLAG-tagged MDA5-2CARD WT and mutants to HA-14-3-3η in transiently transfected HEK293T cells, determined by HA-PD and IB with anti-FLAG and anti-HA. WCLs were probed by IB with anti-FLAG and anti-Actin. **(d)** Binding of FLAG-tagged MDA5 WT and K23R/K43R to HA-tagged LGP2 in transiently transfected HEK293T cells, determined by HA-PD and IB with anti-MDA5 and anti-HA. WCLs were probed by IB with anti-MDA5. Data are representative of at least two independent experiments.

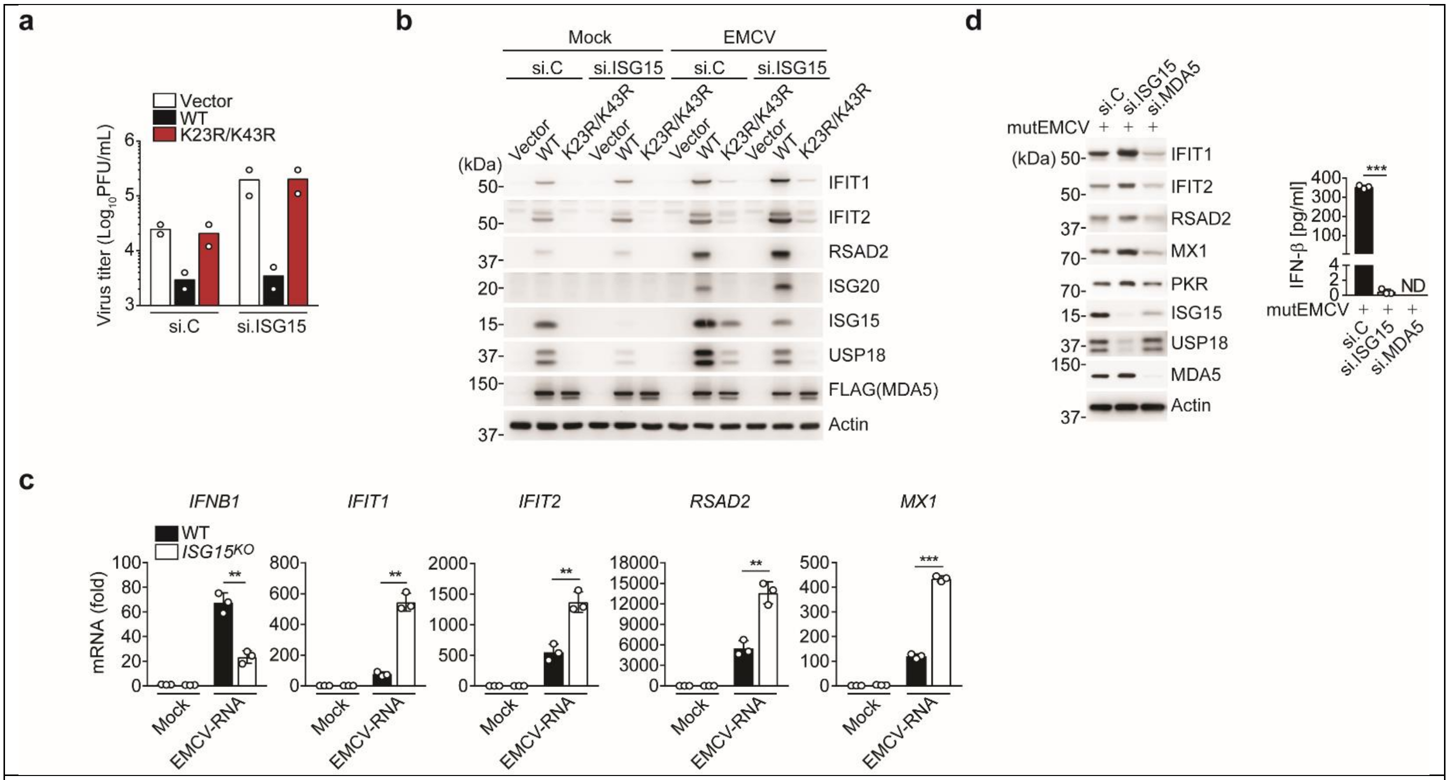


**Extended Data Fig. 5**

**Aberrant ISG upregulation in *ISG15*-deficient cells upon MDA5 stimulation.**

**(a)** EMCV titers in the supernatant of *RIG-I* KO HEK293 cells that were transfected for 24 h with nontargeting control siRNA (si.C) or ISG15-specific siRNA (si.ISG15) and then transfected with either empty vector or FLAG-tagged MDA5 WT or K23R/K43R for 24 h prior to infection with EMCV (MOI 0.001) for 16 h, determined by plaque assay. **(b)** Protein abundance of the indicated ISGs and USP18 in mock-infected or EMCV (MOI 0.001 for 16 h) infected *RIG-I* KO HEK293 cells that were transfected with the indicated siRNAs, and 24 h later, transfected with empty vector or FLAG-MDA5 WT or K23/K43R for 24 h, determined by IB with the indicated antibodies. **(c)** qRT-PCR analysis of *IFNB1* and ISG transcripts in WT and *ISG15* KO HeLa cells that were mock-treated or transfected with EMCV-RNA (0.4 μg/mL) for 16 h. **(d)** Left panel: Protein abundance of the indicated ISGs and USP18 in NHLFs that were transfected for 40 h with the indicated siRNAs and then infected with mutEMCV (MOI 0.1) for 16 h, determined by IB with the indicated antibodies. Right panel: ELISA of IFN-β from supernatants of NHLFs from the same experiment (left panel). Data are representative of at least two independent experiments (mean of $n$ = 2 biological replicates in a; mean ± s.d. of $n$ = 3 biological replicates in c and d). $^{**}p < 0.01$, $^{***}p < 0.001$ (unpaired Student's *t*-test). ND, not detected.

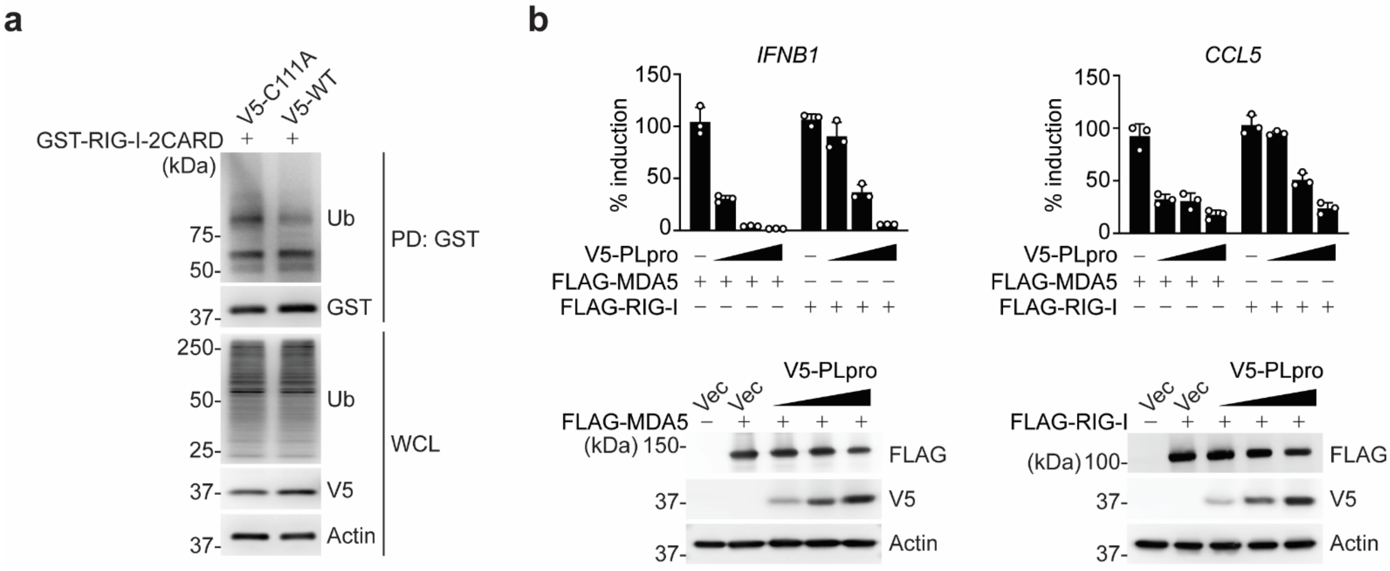


**Extended Data Fig. 6**

**SCoV2 PLpro does not affect RIG-I ubiquitination and preferentially antagonizes the MDA5 pathway.**

**(a)** Ubiquitination of GST-RIG-I-2CARD in HEK293T cells that were transfected with V5-SCoV2 PLpro WT or C111A for 24 h, determined by GST-PD and IB with anti-Ub and anti-GST. WCLs were probed by IB with anti-Ub, anti-V5, and anti-Actin. **(b)** Upper panels: qPCR analysis of *IFNB1* and *CCL5* transcript in HeLa cells that were co-transfected with FLAG-MDA5 or FLAG-RIG-I together with either empty vector (Vec) or increasing amounts of V5-SCoV2 PLpro (10 ng, 25 ng, and 50 ng) for 24 h. Data are presented as percentage of induction relative to the values for cells transfected with the respective RLR (*i.e.* FLAG-MDA5 or FLAG-RIG-I) and vector, set to 100%. Lower panels: WCLs from the same experiment were probed with the indicated antibodies. Data are representative of at least two independent experiments (mean ± s.d. of *n* = 3 biological replicates in b).
